# Hsp40 Affinity to Identify Proteins Destabilized by Cellular Toxicant Exposure

**DOI:** 10.1101/2021.09.22.461285

**Authors:** Guy M. Quanrud, Maureen R. Montoya, Liangyong Mei, Mohammad R. Awad, Joseph C. Genereux

**Affiliations:** Department of Chemistry, University of California, Riverside, CA 92521; Department of Chemistry, Colgate University, Hamilton, NY 13346; Graduate Program of Environmental Toxicology, University of California, Riverside, CA 92521

## Abstract

Environmental toxins and toxicants can damage proteins and threaten cellular proteostasis. Most current methodologies to identify misfolded proteins in cells survey the entire proteome for sites of changed reactivity. We describe and apply a quantitative proteomics methodology to identify destabilized proteins based on their binding to the human Hsp40 chaperone DNAJB8. These protein targets are validated by an orthogonal limited proteolysis assay using parallel reaction monitoring. We find that brief exposure of HEK293T cells to meta-arsenite increases the affinity of two dozen proteins to DNAJB8, including known arsenite-sensitive proteins. In particular, arsenite treatment destabilizes both the pyruvate dehydrogenase complex E1 subunit and several RNA-binding proteins. This platform can be used to explore how environmental toxins impact cellular proteostasis, and to identify the susceptible proteome.

---

Exposure to environmental toxins threatens the structural integrity of proteins.^[1]^ Structural changes due to oxidation, covalent modification, or non-covalent binding can cause proteins to misfold, leading to aggregation or loss of function.^[2]^ For example, the heavy metal arsenic (As) binds to protein sulfhydryl groups and generates reactive oxygen species (ROS).^[3]^ Consequent accumulation of misfolded proteins and oxidative stress activates the Heat Shock Response (HSR) to induce chaperones and restore protein homeostasis.^[4]^ Activation of HSR or other similar misfolded protein stress responses is a common response to heavy metals, electrophilic pesticides/herbicides, and other environmental toxins. While measuring these responses indicates whether a given toxin is likely to be inducing protein misfolding, it does not indicate *which* proteins are misfolding, and hence which cellular pathways are being affected by the exposure.

Most current approaches to identify misfolded proteins measure proteome-wide solvent accessibility by mass spectrometry to infer conformational changes.^[5,6]^ Stability of Proteins from Rates of Oxidation (SPROX) analyzes protein methionine oxidation in cellular lysates, with varying chaotrope concentrations to measure proteins’ ΔG_unfolding_.^[7,8]^ Fast Photochemical Oxidation of Proteins (FPOP) measures the exposure of protein sites in cells or organisms to in situ generated hydroxide radicals.^[9]^ Limited Proteolysis (LiP) measures proteome-wide susceptibility to proteolytic cleavage.^[10]^ Covalent protein painting measures differences in protein folding based on accessible lysine ε-amines after proteins are exposed to electrophilic reagents.^11^ Cellular Thermal Shift Assay (CETSA) measures proteome-wide susceptibility to aggregation with increasing temperature.^12^ Each technique offers a unique approach using quantitative proteomics to assess protein stability in a cell.

Alternatively, the cell identifies misfolded proteins through recognition by chaperones. A highly promiscuous chaperone is Hsp70, which relies upon members of the Hsp40 family to identify and recruit misfolded protein clients for refolding; one third of the proteome relies on this cycle under basal conditions.^[13-16]^ Release of clients from Hsp40 to Hsp70 can be blocked by an H-to-Q mutation in the Hsp40 J-domain, stabilizing misfolded protein binding.^[17]^ DNAJB8 is notable among human Hsp40s for its dual nuclear and cytosolic localization, formation of oligomers, and slow client release kinetics.^[18-21]^ We previously used affinity purification and quantitative proteomics to identify hundreds of cellular protein clients of overexpressed human Hsp40 DNAJB8^H31Q^ with high reproducibility and statistical confidence.^[22]^ Herein we exploit the ability of DNAJB8^H31Q^ to recognize misfolded protein clients to develop a platform for identifying proteins that are destabilized in response to exogenous stress (**Figure 1**). We demonstrate this approach in HEK293T cells treated with trivalent arsenic, a toxic metal that causes widespread damage to nucleic acids and proteins, leading to genomic and metabolic instability.^[23]^

**Figure 1.**
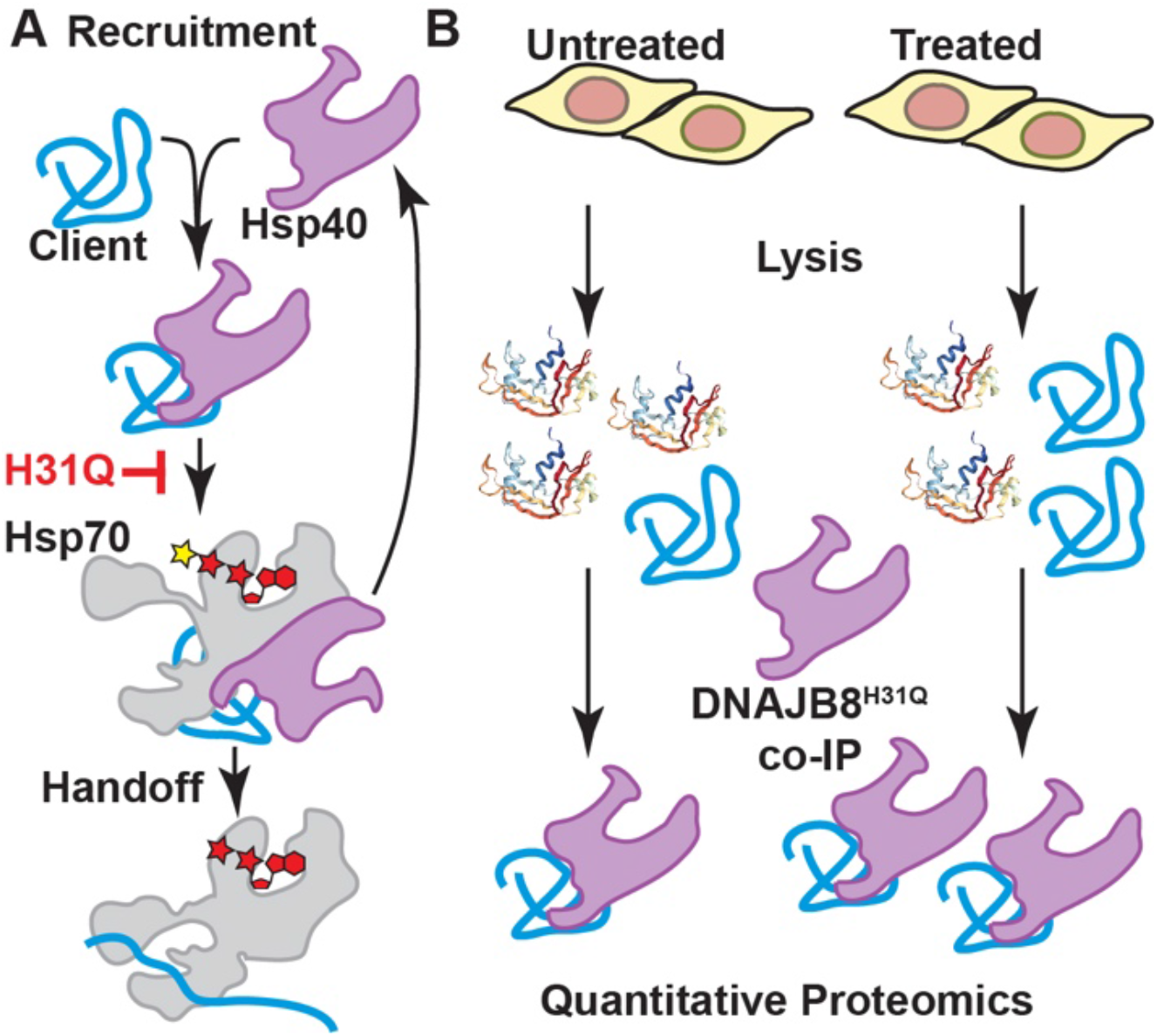
Design of the Hsp40 affinity assay for misfolded proteins. A) DNAJB8 binding and handoff to Hsp70 is interrupted by an H31Q mutation in the J-domain. B) If cellular treatment increases the misfolded population of a DNAJB8 client protein, then the apparent affinity of that protein for DNAJB8^H31Q^ will increase.

Protein misfolding stresses leads to extensive transcriptional, translational, and post-translational remodeling of the cell.^[24]^ To isolate stress-induced protein misfolding from these pleiotropic effects, we limited cellular exposure to brief 15 min treatments. We validated that 15 min. 500 µM sodium meta-arsenite (NaAsO_2_) induces expression of the HSR target HSPA1A in HEK293T cells (**Figure S1**). Because HSR is activated by misfolded protein accumulation, this suggests that 500 µM arsenite treatment causes protein misfolding in only 15 min. This response is not suppressed by overexpression of ^Flag^DNAJB8^H31Q^ (**Figure S1**), indicating that our recognition element for misfolded protein does not prevent protein destabilization.

We used the experimental approach illustrated in **Figure 2A** to determine proteins that misfold in response to arsenite exposure. ^Flag^DNAJB8^H31Q^ was transiently overexpressed in HEK293T cells, followed by 15 min. NaAsO_2_ treatment and immediate Flag immunoprecipitation from cellular lysate. Co-immunoprecipitated proteins were identified and quantified by LC/LC-MS/MS in concert with TMT isobaric tagging.^[25]^ Overall, 24 biological replicates (12 treated and 12 controls) were analyzed through four 6-plex TMT runs. Most observed protein clients slightly increase affinity to DNAJB8 following arsenite treatment, likely because peptides from proteins with increased DNAJB8 binding following arsenite treatment are more represented in the pooled peptide mixture, and thus have higher chances of identification during shotgun proteomics. However, the bulk DNAJB8 associated proteome does not change (**Figure 2B**). 24 proteins have significantly greater affinity for ^Flag^DNAJB8^H31Q^ in response to the arsenite treatment. These proteins include PDHA1 and 17 ribosomal RNA-binding proteins, including HNRNPA0, TDP-43, RACK1, and RPS16 (**Figure 2C** and **Table S1**). Arsenite generally induces misfolding and aggregation of RNA-binding proteins into stress granules.^[27,28]^ However, canonical stress granule markers^[29]^ G3BP1 and eIF4G1 are not enriched in DNAJB8^H31Q^ pull-downs from arsenite-treated cells, indicating that DNAJB8^H31Q^ is not co-precipitating intact stress granules (**Table S1**). TDP-43 is a ribonuclear protein that forms aggregates in ALS and other proteinopathies.^[30]^ It accumulates in cytoplasmic and nuclear condensates in response to arsenite treatment, due to displacement from RNA and post-translational modification.^[31]^ Yeast RACK1 migrates to stress granules in response to arsenite.^[32]^ It is particularly interesting that PDHA1, the alpha subunit of the pyruvate dehydrogenase complex (PDC),^[33]^ is destabilized by arsenite. Inhibition of PDC is a major contributor to arsenite-induced metabolic disfunction.^[34]^ It is encouraging that our assay primarily finds proteins that are known to be arsenite-sensitive.

**Figure 2.**
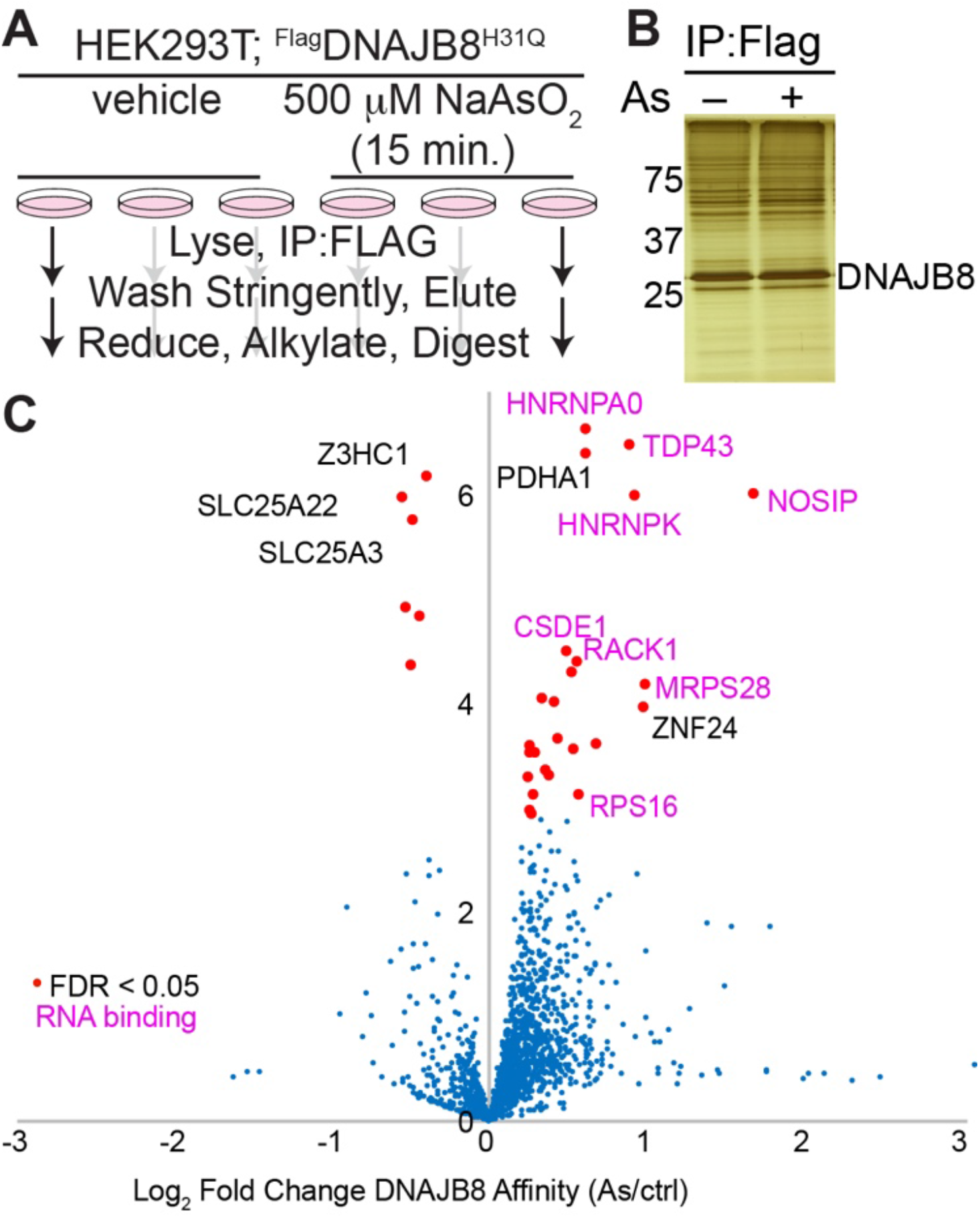
Arsenite treatment increases the affinity of a select subset of proteins with DNAJB8^H31Q^. A) Experimental protocol. B) Representative silver stain for proteins co-immunoprecipitated with DNAJB8H31Q. C) Volcano plot illustrating the effect of cellular As treatment on protein interactions with DNAJB8^H31Q^. Red dots represent proteins with significantly increased interaction with DNAJB8^H31Q^, using a false discovery rate threshold (FDR) of 5% (n = 12 biological replicates in 4 TMT-AP-MS runs). Protein names in purple are RNA-binding proteins.

To determine whether the arsenite response is specific, we treated cells with Cd^2+^, another heavy metal that induces cellular apoptosis through generation of ROS.^[35]^ 15 min. treatment with 200 µM Cd(NO_3_)_2_ induces HSR in HEK293T cells (**Figure S2**). Cells were treated with Cd or vehicle for 15 min. and then immediately lysed, and the DNAJB8^H31Q^ interacting proteome quantified by TMT-AP-MS (**Figure 3** and **Table S2**). Unlike with arsenite, the only proteins destabilized by acute Cd treatment are PDHA1 and BAZ1B. Rather, many proteins slightly lose affinity for DNAJB8 following treatment, perhaps due to direct binding to Cd. As with arsenite, Cd exposure inhibits pyruvate dehydrogenase activity. TDP-43 association with DNAJB8 is unchanged, despite reports that Cd^2+^ treatment (100 µM, 2 h) promotes TDP-43 aggregation in Cos7 cells.^[30]^ The difference in proteome destabilization between the two heavy metals suggests that ROS generation is not adequate to explain their effects on protein stability following acute arsenite exposure.

**Figure 3.**
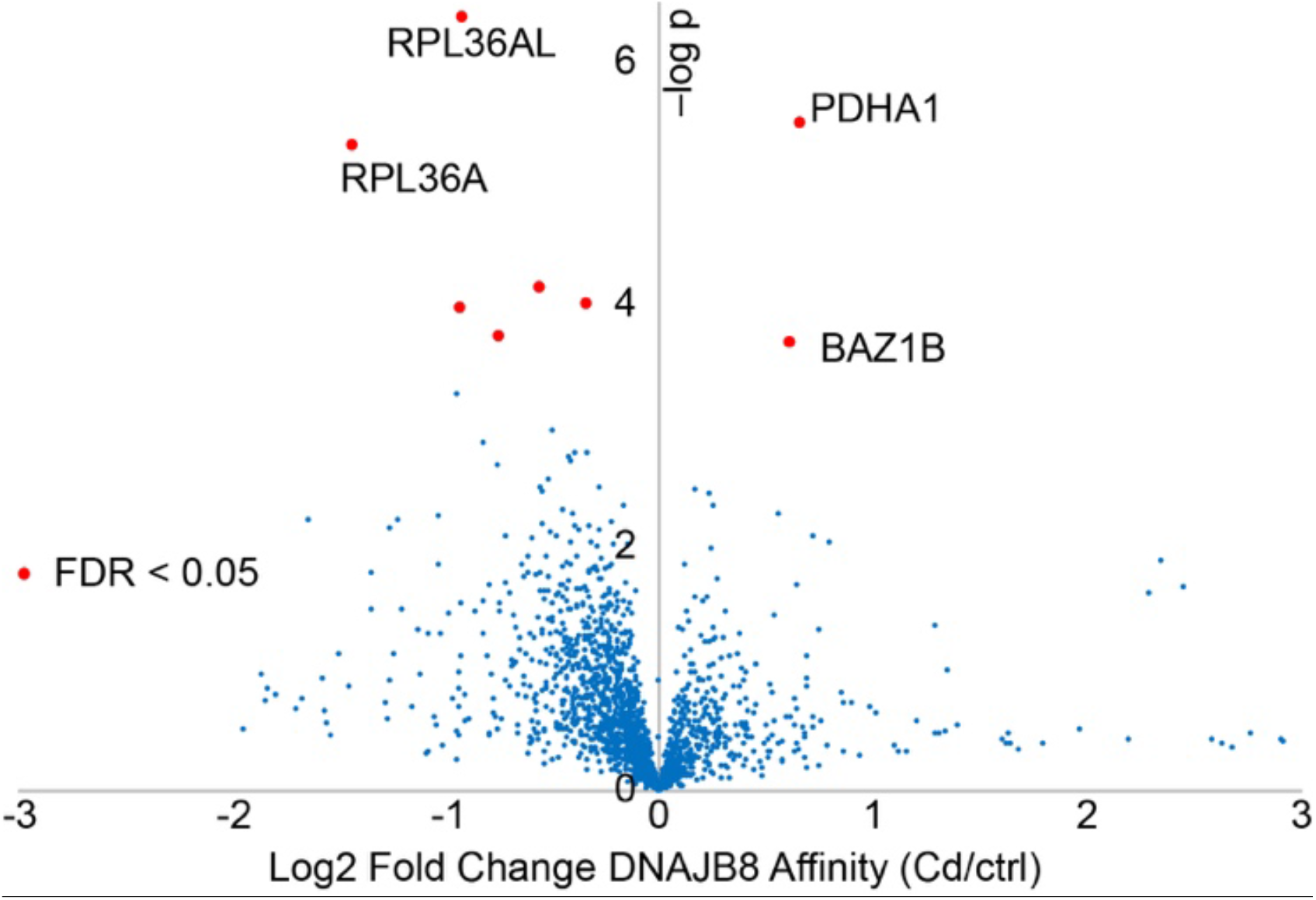
Cadmium treatment does not affect the affinity of many proteins for DNAJB^8H31Q^. The experimental protocol is similar to in **Figure 1A**, except that HEK293T cells were treated with 200 *µ*M CdCl2 for 15 min. prior to lysis. PDHA1 is the only protein below the false discovery rate threshold of 0.05 (from n = 12 biological replicates in 4 TMT-AP-MS runs).

Although our short treatments and immediate processing avoid many pleiotropic effects (e.g. altered transcription, translation, trafficking, and degradation, etc.), unforeseen variables besides protein stability could impact DNAJB8-client affinity. Hence we applied an orthogonal assay, LiP, to validate and prioritize proteins with arsenite-induced binding to DNAJB8 (**Figure 4A**). LiP as a discovery technology is challenged by the need to identify and quantify peptides in a proteome that has been rendered complex by the use of two orthogonal proteases; for example, hit overlap with SPROX from the same samples is about ∼20%.^[36]^ We targeted select peptides from our hit proteins to alleviate this challenge. We treated cells with arsenite or vehicle for 15 min., followed by immediate lysis. Lysates were treated with varying concentration of proteinase K for 1 min, heat-quenched, tryptically digested, and peptides from candidate proteins quantified by Parallel Reaction Monitoring (PRM).^[37]^ If a cellular treatment increases the PK proteolysis yield of a peptide, that implies that the protein is destabilized in the vicinity of that sequence.^[11]^ Limited proteolysis is a local measure of protein conformation and thus is subject to false negatives at the protein level. Hence, a given peptide being equally sensitive to proteolysis with or without cellular treatment does not imply that the entire protein is unaffected.

**Figure 4.**
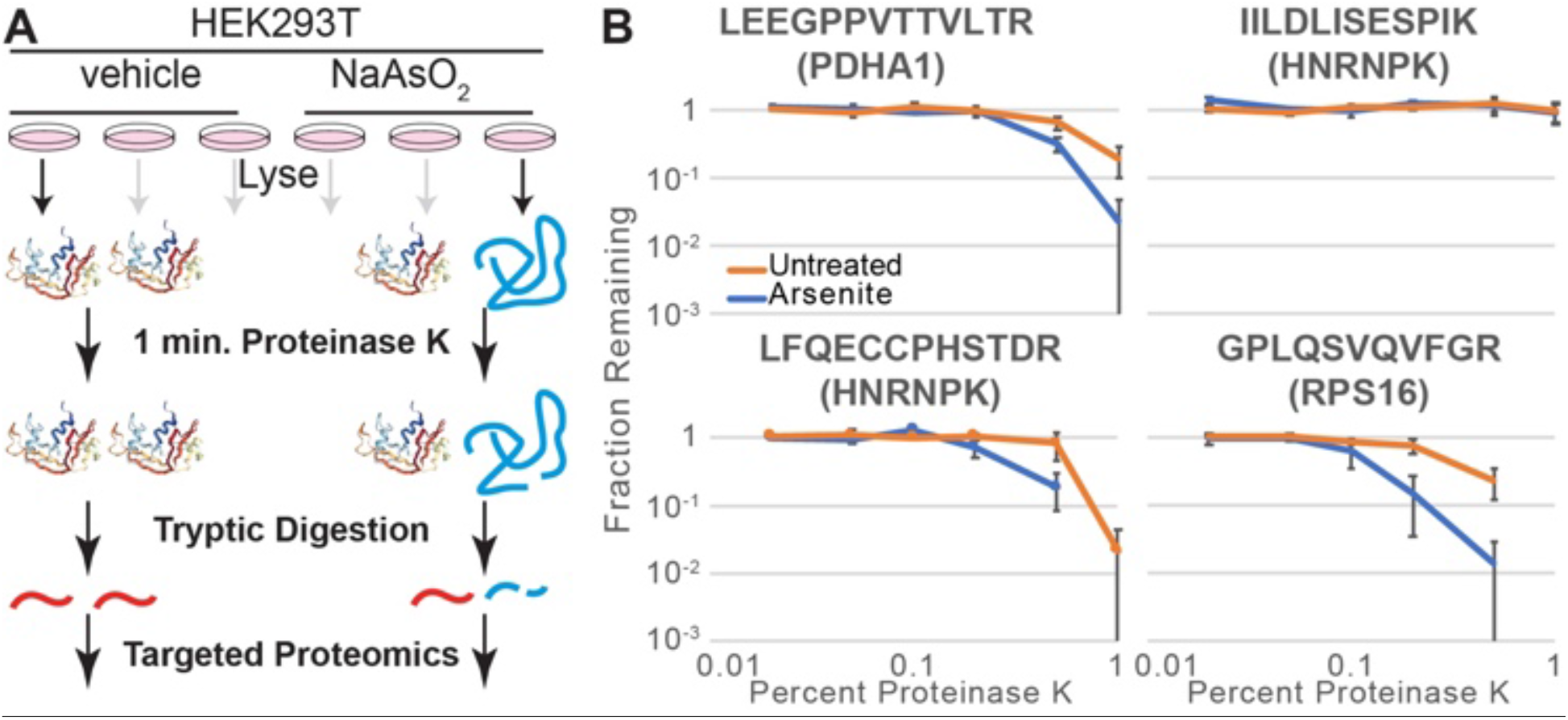
A) Schematic illustrating how limited proteolysis differentiates between different conformations of a protein. B) Proteinase K susceptibility curves for four peptides as monitored by LiP-PRM. Error bars represent standard error (n = 3).

We determined the proteolytic susceptibility with and without 15 min. cellular arsenite treatment for 13 peptides from PDHA1, RSP16, RPS3, TDP-43, HNRNPK, RACK1, and HNRNPA0. Although RPS3 was a lower significance protein from our screen, we included it in these targeted LiP experiments because arsenite causes release of RPS3 from RNA,^[38]^ which could affect protein stability. Peptides from PDHA1 and RPS16 are more susceptible to proteolysis upon arsenite treatment, implying that arsenite destabilizes these proteins (**Figures 4B** and **S3A**). The peptides chosen for TDP-43, HNRNPA0, RACK1 and RPS3 do not show a clear increase in proteolytic sensitivity; those proteins might still be destabilized by arsenite in unsampled regions of the protein. HNRNPK increases sensitivity at two peptides, but not at three others. No peptide became less sensitive to PK following cellular arsenite treatment. We expanded the TDP-43 evaluation to four additional peptides, of which three showed arsenic sensitivity. Surprisingly, all three peptides come from structured regions within the RNA-binding domains of the protein (**Figure S3B**).^[39]^ The two peptides that do not have increased proteolytic susceptibility in response to cellular arsenic exposure are in intrinsically disordered regions; one is in the linker between the RNA binding domains and the other is in the unstructured C-terminus. The general validation of protein destabilization by an orthogonal method demonstrates that DNAJB8^H31Q^ affinity successfully identified proteins that are destabilized by arsenite treatment.

The sensitivity of PDHA1 to arsenite is surprising. PDC inhibition by arsenite is generally ascribed to arsenite coordination to vicinal thiols in the lipoamide cofactor,^[3]^ though other evidence strongly points to lipoamide binding being unneccesary for inhibition by arsenic.^[40]^ PDC is composed of three subunits, including an E1 heterotetramer containing two PDHB and two PDHA1.^[33]^ Lipoamide is anchored covalently to DLAT in the E2 subunit, but reacts in the groove of PDHB in the E1 subunit. We considered that arsenite destabilizes the rest of the E1 subunit. PDHB did show increased affinity to DNAJB8^H31Q^ in our initial screen, but with low significance (FC = 1.3 ± 0.3, q = 0.06). DLAT was observed to not significantly bind to DNAJB8^H31Q^ and unaffected after arsenite exposure (FC = 0.93 ± 0.48, q = 0.69). Because LiP measures local proteolytic sensitivity,^[41]^ we used targeted LiP to determine whether PDHA1 and PDHB are globally destabilized. Eleven peptides from PDHA1 and PDHB were selected, covering most of the protein sequences (**Figure S3C**). Most locations on both proteins are destabilized by cellular arsenite treatment (**Figure S4**), indicating that the proteins undergo an extensive conformational change with arsenite treatment. Despite destabilization throughout the E1 subunit, this complex remains intact following acute arsenite treatment (**Figures S5-S6** and **Supplemental Discussion**).

In conclusion, we have developed a platform to identify proteins that are destabilized in response to cellular stress. The DNAJB8^H31Q^ immunoprecipitation assay identifies destabilized proteins using the same criterion as the cell: increased binding to a chaperone. Identification of likely destabilized proteins in the screening step enables targeted LiP-PRM as a mechanistically orthogonal, and technically straightforward, validation step. Using this technology, we identified proteins that are destabilized by just a brief 15 min. cellular arsenite exposure. Most of these proteins are known to be functionally perturbed by arsenite, with changes in post-translational modifications and even aggregation. However, it has not previously been demonstrated that these proteins are destabilized, nor that arsenite can induce conformational changes inside living cells within 15 min. of treatment. Furthermore, we have found that arsenite destabilizes both members of the E1 subunit of PDC. This platform will be useful in further understanding how environmental toxins and toxicants perturb proteome integrity.

## Supporting information

Supplemental Tables 3 through 8

## Acknowledgements

We thank H.T. Wu for assistance with Skyline. This work was supported by the University of California, Riverside and a Society of Analytical Chemists of Pittsburgh Starter Grant.

