## Supplemental Text and Figures for "Hsp40 Affinity to Identify Proteins Destabilized by Cellular Toxicant Exposure"

*and Joseph C. Genereux**

**Page Contents**

| Page | Contents |
|---|---|
| S1 | **Table of Contents** |
| S2 | **Materials** |
| S3 | **Supplemental Methods** |
| S14 | **Supplemental Results and Discussion** |
| S19 | **Supplemental Figures** |
| S48 | **Supplemental References** |

**Supplemental Table 1** is provided as an external file.

**Supplemental Table 2** is provided as an external file

**Supplemental Tables 3** through **8** are provided as an external file.

**MATERIALS**

Bovine Serum Albumin (BSA), Dulbecco's Modified Eagle Media (DMEM), Dublecco's phosphate-buffered saline (DPBS), 10 cm plates, and 6 well plates were from VWR. Roche Protease Inhibitor cocktail w/o EDTA (PIC), 1,4-Dithiothreitol (DTT), 4-(2-hydroxyethyl)-1-piperazineethanesulfonic acid (HEPES), sodium meta arsenite ($NaAsO_2$), cadmium nitrate tetrhydrate ($Cd(NO_3)_2$) were from Sigma Aldrich. Sodium chloride (NaCl), Tris-Hydrochloride (Tris-HCl), Triton X-100, sodium deoxycholate ($C_{24}H_{40}O_4$), Potassium chloride (KCl), Magnesium chloride ($MgCl_2$), calcium chloride ($CaCl_2$), Silver Nitrate ($Ag(NO_3)_2$), Sodium thiosulfate ($Na_2S_2O_3$), urea, calcium acetate ($Ca(O_2C_2H_3)_2$), glycerol, pierce trypsin protease, sodium dodecyl sulfate (SDS), poly D-lysine, sequence grade trypsin were from Thermo Fisher Scientific. Proteinase K (PK) was from Promega. Nanopure water was purified using a Millipore Milli-Q Laboratory lab 4 Chassis Reagent Water System. 5 µm Aqua C18 resin and 3 µm Aqua C18 Resin were from Phenomenex. Sepharose-4-B beads, anti-M2 Flag beads, tris (2-carboxyethyl)phosphine hydrochloride (TCEP), and iodoacetamide were from Millipore Sigma. 250 µm diameter fused silica columns and 100 µm diameter fused silica columns were from Agilent. Strong cation exchange resin was from Partisphere, GE Healthcare. Rapigest was from Aobious (Gloucester, MA). TMT-6plex isotopic labels were from Pierce. Bradford reagent was purchased from Bio-rad. All other antibodies were obtained from Proteintech.

**SUPPLEMENTAL METHODS**

*Cell Culture:* HEK239T cells were obtained from the ATCC and maintained in DMEM with 10% FBS. Cell transfection was performed by the calcium phosphate method. Briefly, 5 μg of DNA in 1 mL 250 μM $CaCl_2$ is vortexed while adding dropwise 1 mL HBS 2X for 10 seconds at ambient temperature, the transfection solution is promptly (≤ 15 min) added dropwise to cells at about 50% confluency, and the cell media is changed between 12 and 16 h. A positive transfection control is performed with GFP alongside each transfection. These transfection amounts were used for 10 cm dishes; proportional amounts of reagents were used for transfection of different sized plates.

Immunoprecipitation/TMT-MudPIT Analysis of As and Cd-Interactomes

*Immunoprecipitation of $^{Flag}$DNAJB$^{H31Q}$:* DNAJB8 TMT-AP-MS experiments were performed as previously described[1]. For each sixplex TMT-AP-MS, six 10 cm plates of HEK293T cells were transfected by the calcium phosphate method with 5 μg of plasmid DNA encoding $^{Flag}$DNAJB8$^{H31Q}$ in the pFLAG backbone[1]. Plates were treated with heavy metal salts or vehicle at 40-46 hours post transfection. Cells were harvested by scraping in DPBS and lysed in 9 parts RIPA Buffer (150 mM NaCl, 50 mM Tris pH 7.5, 1% Triton X-100, 0.5% sodium deoxycholate, 0.1% SDS) and 1 part 10x PIC for 30 min on ice. Lysate was separated from cell debris by centrifugation at 21,000 *x g* for 15 minutes at 4 °C. Protein in the lysate was quantified by Bradford. Lysates were pre-cleared with 15 μL Sepharose-4B beads for 30 min at 4 °C, then centrifuged at 1,500 *x g* for 1 min to pellet beads. Lysate was then separated and incubated with 15 μL of M2 anti-Flag Magnetic Beads and rotated overnight at 4 °C. The anti-Flag beads were washed the next day four times with RIPA buffer. Each wash included rotation for 10 minutes at ambient temperature. Proteins bound to the anti-Flag beads were eluted by boiling for 5 min at 100 °C in 30 μL of Laemmli concentrate (120 mM Tris pH 6.8, 60% glycerol, 12% SDS, brilliant phenol blue to color). 5 μL of the elutes were saved for silver stain analysis and the remainder was prepped for mass spectrometry.

*Silver Stain.* Silver stain was used to assess the amount of the $^{Flag}$DNAJB8$^{H31Q}$ in eluates after the immunoprecipitation and potential differences in the co-immunoprecipitated protein levels between conditions[2]. DTT was added to a final concentration of 170 mM, and eluates boiled for 5 min at 100 °C prior to SDS-PAGE separation. Gels were fixed overnight in 30% ethanol/10% acetic acid or for a few hours with 50% methanol/12% acetic acid. Gels were washed in 35% ethanol three times for 20 minutes each, sensitized for 2 minutes (0.02% $Na_2S_2O_3$ in $H_2O$), washed three times for 1 minute each in $H_2O$, and stained for 30 minutes to overnight in Ag staining solution (0.2% $AgNO_3$, 0.076% formalin). After two one minute rinses in $H_2O$, gels were developed with 6% $NaCO_3$/0.05% formalin/0.0004% $Na_2S_2O_3$. Development was stopped with 5% acetic acid. Gels were imaged on a white-light transilluminator (UVP).

*TMT-MudPit.* Eluates reserved for mass spec analysis were prepared for TMT-AP-MS. Only MS quality organic solvents were used during sample preparation. The composition for buffer A is 5% acetonitrile, 0.1% formic acid in water. The composition for Buffer B is 80% acetonitrile, 0.1% formic acid. The composition for Buffer C is 500 mM ammonium acetate in Buffer A. Proteins in eluates were precipitated by methanol/chloroform precipitation. Pellets were then air-dried and resuspended in 1% Rapigest in water. Resuspended protein solutions were then diluted to 50 µL in 100 mM HEPES, pH 8.0, and reduced with 10 mM TCEP for 30 min at 37 °C. Protein solutions were then alkylated with 5 mM iodoacetamide for 30 min in the dark at ambient temperature. 0.5 µg sequencing grade trypsin was added to the protein solution for digestion overnight at 37 °C with agitation (600 rpm). TMT isotopic labels were resuspended (100 ug/80 µL acetonitrile) and 40 µL of label was added to each 60 µL sample of digested peptides. Low excess of TMT to peptide has been shown to minimize side-product generation during labeling[3]. Samples were labeled for 1 h at ambient temperature. Labeling was quenched with 0.4% ammonium bicarbonate at ambient temperature for 1 h. Samples were pooled, acidified, centrifuged for 30 min at 21,100 *x g* to remove any insoluble debris. Samples were then dried by centrifugal evaporation to 10 μL. Solutions were then brought to 200 μL in Buffer A, incubated at 37 °C for 1 h, and centrifuged for 30 minutes at 21,100 *x g*. Solution was transferred to new low-binding tubes (Eppendorf) and the process of heat-spinning was repeated three more times to complete elimination of Rapigest.

There were two different set ups used for mass spec analysis. The first arsenic exposure TMT-MS run was performed as one-dimensional LC/MS/MS on an Orbitrap Fusion Tribrid mass spectrometer (Thermo) interfaced with a nanoAquity UPLC (Waters) system. Samples were loaded onto a single phase loading column. The loading column was prepared by polymerizing a Kasil 1624 frit into a 250 μm diameter fused silica capillary and then packed with 2.5 cm reversed-phase 5 μm Aqua C18 resin. Analytical columns were prepared by pulling 100 μm diameter fused silica columns with a P-2000 laser tip puller (Sutter Instrument Co., Novato, CA), followed by packing with at least 20 cm reversed-phase 3 μm Aqua C18 resin. LC separation was achieved with a gradient from water to acetonitrile (ACN). Both LC buffers contained 0.1% formic acid. The percent amount of ACN increased from 0% to 9% ACN for the first 10 minutes. The amount of ACN increased to 30% for the next 110 minutes. The percent amount of ACN increased to 90% ACN for 10 minutes and then remained at 90% for 20 more minutes. The percent amount of ACN decreased to 9% ACN over 1 minute and remained at 9% ACN for the final 10 minutes. The flow rate was 300 nL/min. Eluted peptides were ionized by electrospray (2.8 kV) and scanned from 400 to 1500 m/z in the orbitap with resolution 120,000 in data dependent acquisition mode. The top ten peaks with charge states of 2+, 3+, 4+, 5+, 6+, 7+ from each full scan were fragmented by HCD using a stepped collision energy of 37.6%, 40%, and 42.4%, a 100 milliseconds activation time, and a resolution of 30,000. Dynamic exclusion parameters included excluding after 1 times, 5 seconds exclusion duration, and mass tolerance of 10 ppm for high and low. Peptides were selected with 0.7 Da isolation window in the quadrupole.

All other MS runs were performed with a two-dimensional LC/MS/MS setup on an LTQ Orbitrap Velos Pro hybrid mass spectrometer (Thermo) interfaced with an Easy-nLC 1000 (Thermo) according to standard MuDPIT protocols[4]. Samples were loaded onto a triphasic loading column for analysis. Triphasic loading columns were prepared by polymerizing a Kasil 1624 frit into a 250 µm diameter fused silica capillary. The column was then packed with 2.5 cm of reversed-phase 5 µm Aqua C18 resin, followed by 2.5 cm of 5 µm strong cation exchange resin, and again with 2.5 cm of reversed-phase 5 µm Aqua C18 resin. Analytical columns were prepared by pulling 100 µm diameter fused silica columns with a P-2000 laser tip puller (Sutter Instrument Co., Novato, CA), followed by packing with at least 20 cm reversed-phase 3 µm Aqua C18 resin. Analysis was performed using a 11-cycle chromatographic run, with progressively increasing concentrations of ammonium acetate salt bumps injected prior to each cycle (0% C, 10% C, 20% C, 30% C, 40% C, 50% C, 60% C, 70% C, 80% C, 100% C, 90% C+ 10% B; balance of each buffer A), followed by acetonitrile gradient (5 min from 1% B to 7% B, 60 min to 55% B, 15 min to 100% B, 5 min at 100% B, 5 min to 1% B; 300 nL/min flow rate). Eluted peptides were ionized by electrospray (3.0 kV) and scanned from 110 to 2000 m/z in the Orbitrap with resolution 30,000 in data dependent acquisition mode. The top ten peaks with charge states of 2+, 3+, or 4+ from each full scan were fragmented by HCD using a stepped collision energy of 36%, 42%, and 48%, a 100 milliseconds activation time, and a resolution of 7500. Dynamic exclusion parameters were 1 repeat count, 30 milliseconds repeat duration, 500 exclusion list size, 120 seconds exclusion duration, and 2.00 Da exclusion width.

For each run, MS/MS spectra were extracted using MSConvert (version 3.0.21144-1f7ddf52b) with Peak Picking Filtering. MS/MS spectra was then searched by Fragpipe against a Uniprot human proteome database (06/11/2021 release) containing 40858 human sequences (longest entry for each protein). MS/MS spectra were also searched against 20429 select decoys (e.g albumen, porcine trypsin, contaminats etc.). FragPipe searches allowed for static modification of cysteine residues (57.02146 Da, acetylation), and N-termini and lysine residues (229.1629 Da, TMT-tagging), half tryptic peptidolysis specificity, and mass tolerance of 20 ppm for precursor mass and 20 ppm for product ion masses. Spectra matches were assembled and filtered by MSFragger (version 3.2). Decoy proteins, common contaminants, immunoglobulins and keratins were filtered from the final protein list. Quantitation in FragPipe was performed by averaging TMT reporter ion intensities for all spectra associated with an individual peptide.

The Mass Spectrometry Proteomics Data was deposited to the ProteomeXchange Consortium by using PRIDE[5] Submission tool. The Data set identifier is PXD028110. Reviewer Information is Username: and Password: P9Us0u5s

Statistical Analysis:

Protein-level intensities were normalized to the intensity of bait (DNAJB8) in each TMT channel. To combine the multiple TMT runs, we used a version of the scaled reference approach[6]. A scaling factor was obtained from averaging the bait-normalized integrated TMT reporter ion intensities for each protein across the 3 control conditions in each AP-MS run (**Supp Table 1**). Each bait-normalized protein intensity was then divided by this scaling factor. Unadjusted p-values were converted to q-values (local false discovery rates) using Storey's modification to the method of Benjamini and Hochberg as follows[7,8,9,10,11]. Unadjusted p-values were ranked in increasing order and the q-value ($q_{BH}$) for the *i*th protein determined from

$$q_{BH,i} \quad = min_{i \leq j \leq n} \frac{pn}{i}$$

where *n* represents the number of quantified proteins. Storey's modification is performed by determining the overrepresentation of low p-values to infer a global false discovery rate, and then scaling local false discovery rates accordingly[12,13]. The $\pi$-factor for this scaling was calculated to be 0.84 for the arsenic treatment TMT-AP-MS data set and 0.79 for the cadmium treatment TMT-AP-MS set.

Limited Proteolysis

*Arsenite Treatment and LiP*: Six 10 cm plates seeded with HEK293T cells were transfected with 5 µg of DNA encoding $^{Flag}$DNAJB8$^{H31Q}$ by the calcium phosphate method as described above. Three plates were incubated with 500 µm $Na_3AsO_3$ (As) and 3 plates were incubated with water for 15 min at 37 °C, 40-46 hours post transfection. Plates were immediately harvested by scraping in DPBS and lysed with 9:1 Native Lysis Buffer (20 mM HEPES, 150 mM KCl, 10 mM $MgCl_2$, 0.3% Triton, pH 7.5): 10x PIC for 30 min in ice. Lysates were separated from cell debris by centrifugation at 21,000 *x g* for 15 min at 4 °C. 200 µg aliquots were prepared for limited proteolysis after Bradford protein quantification.

Limited Proteolysis procedure was optimized from standard protocols[14]. 1 mg/ml stocks were made from 25 mg of lyophilized Proteinase K (PK) dissolved in a storage buffer (50 mM Tris-HCl, 2 mM calcium acetate, pH 8.0) suitable for PK and stored at –70 °C. The following concentrations of PK were prepared from serial dilutions from 1mg/ml aliquot: 0.5 mg/ml, 0.2 mg/ml, 0.1 mg/ml, 0.05 mg/ml, 0.02 mg/ml, and added to lysate to yield 1:200, 1:500, 1:1000, 1:2000, and 1:5000 wt/wt protease: substrate protein ratios respectively. For each digestion, 2 µl PK was added to a 200 µg aliquot of protein lysate and incubated for 1 min at 25.0 °C. Samples were then boiled for 5 min to quench PK activity. For the method analyzing five TDP-43 peptides, we did not perform the 1:5000 wt/wt condition. Three separate digestions were performed for the no PK condition for each lysate sample. The sample set-up and the calculation of fraction remaining for each peptide is shown in **Schematic S1**:

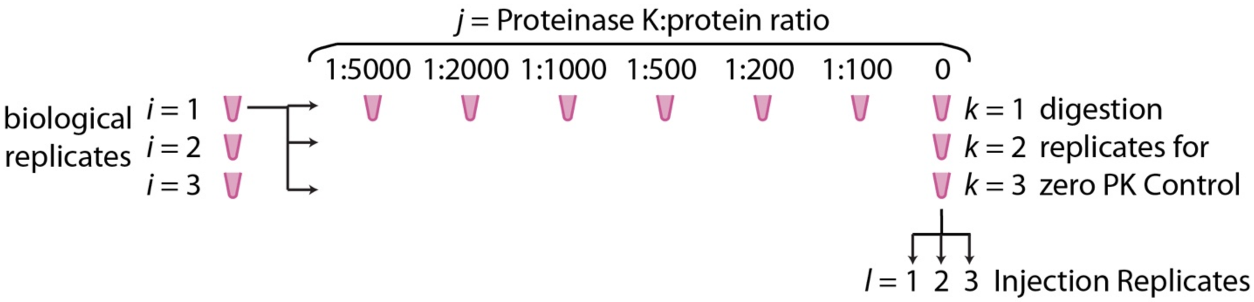


$$\text{fraction remaining } (i) = \frac{\frac{1}{3}\sum_{l=1}^{3}(I_{i,l})}{\frac{1}{9}\sum_{k=1}^{3}\sum_{l=1}^{3}(I_{i,0,k,l})}$$

**Schematic S1**

*Mass Spec Prep*: Methanol/chloroform precipitation was performed on each sample following LiP. Pellets were solubilized in 8 M urea in 50 mM Tris pH 8 and reduced with 10 mM TCEP in 50 mM Tris pH 8 for 30 min at ambient temperature. Samples were then alkylated with 5 mM iodoacetamide for 30 min in the dark at ambient temperature. Solutions were then diluted with 50 mM Tris pH 8 to 2 M urea and brought to 10 mM $CaCl_2$ to improve protease activity. Sequencing grade trypsin was added at 100:1 substrate: enzyme ratio. Solutions were agitated at 600 rpm for 20 hours at 37 °C. Trypsin digestion was quenched by adding formic acid to at least 5% of solution and until the pH of the solution is measured at 2 by litmus paper. Internal standard $NH_2$-VFFAEDVGSNK-$CO_2H$ was spiked into the sample from a 10 mM stock. The final concentration of internal standard in the protein sample was 83 nM. The Protein sample was diluted with Buffer A to 1 mg/ml. From each lysate sample, 18 samples were prepared: 3 each of the Trypsin control (No PK), 1:5000 PK:protein, 1:2000 PK:protein, 1:1000 PK:protein, 1:500 PK:protein, 1:200 PK:protein, and 1:100 PK:protein. Three biological replicates were prepared for As and for untreated, yielding a total of 108 samples.

*PRM selection and analysis*: Thirteen peptides were selected from PDHA1, TDP43, RACK1, HNRNPA0, RPS16, HNRNPK, and RPS3 for the initial LiP screen experiment. These proteins were chosen as having significantly increased interaction with DNAJB8 in response to arsenite (see **Supp. Table 1**). Our initial peptide identification searches, using a different software than Fragpipe, did not identify NOSIP, CSDE1, MRPS28, and so we did not include these proteins in our LiP survey. Retention times for each peptide were determined by running unscheduled PRM runs from a trypsin digested lysate. Picky software allowed peptides to be chosen for PRM analysis[15]. Eleven peptides from PDHA1 and PDHB were selected for the E1 subunit LiP, and five peptides from TDP43 were chosen for the TDP43 LiP experiment. Fragmentation patterns for all peptides were uploaded into Skyline software for analysis[16].

Samples were analyzed using two dimensional LC/MS/MS on an LTQ Orbitrap Velos hybrid mass spectrometer (Thermo) interfaced with an Easy-nLC 1000 (Thermo). 6 µg of protein were injected in each run onto a loading column packed with 2.5 cm of 5um Aqua C18 resin and washed prior to separation on the analytical column. Analytical columns were prepared by pulling 100 µm diameter fused silica columns with a P-2000 laser tip puller (Sutter Instrument Co., Novato, CA), followed by packing of 23 cm of reversed-phased 3 µm Aqua C18 Resin. Peptides injected were scanned over scheduled 10 min windows centered around average retention time, and isolated with a 2.0 m/z isolation window. Peptides were fragmented with CID with a normalized energy of 35, activation Q of 0.25 and activation time of 10 msec. MS2 were acquired in the Orbitrap at 7500 resolving power and saved in profile mode. Peptide separations by LC-MS proceeded between Buffer A (5% acetonitrile:95 % water: 0.1% formic acid) and Buffer B (80% acetonitrile: 20% water: 0.1% formic acid) over a 100 min gradient with the following segments: 1-5 min: 1-6% Buffer B. 5-75 min: 6-29% Buffer B. 75-80 min: 29-100% Buffer B. 80-85 min: 100% Buffer B. 85-90 min: 100-1% Buffer B. 90-100 min: 1% Buffer B. Flow rate was 300 nl/min.

*Limited Proteolysis: Data Analysis*: Three technical runs were run for each biological replicate, except only a single technical replicate was ran when using the method analyzing five TDP-43 peptides. For each control (no PK) run, three separate digestions were prepared. Integrated fragment intensities were calculated in Skyline. Integrated fragment intensities were normalized to the integrated fragment intensity of the internal standard for method optimization. However, we did not use internal standard normalization for the digested samples, as we found unacceptable interference in the digested samples for this peptide. The integrated fragment intensities for each set of three technical replicates are averaged and normalized to the averages of the three no-PK controls (which themselves were run in technical triplicate) (**Schematic S1**). Proteolytic susceptibility curves were made from graphs that plotted relative fragment intensity against increasing PK concentration. Differences between curves are assessed based on the summed fractions remaining across data points from 2000:1 to 100:1, and the significance of these sums determined by one-tailed Student's t test (**Supp Table 3**). Coefficient of Variation (CV) were calculated from ten technical replicates of a Trypsin control and CV of biological replicates were calculated among three biological replicates for both LiP experiments. We should note that because our analysis does not assume linearity between integrated peptide response and actual peptide levels, no effort was made to establish whether the peptides observed are in the linear quantitative regime[17].

Assessing HSF1 Activation after Applied Metal Stress

Two 10 cm plates seeded with HEK293T cells were transfected at 40-60% confluency. One plate was transfected with 5 µg of DNA encoding $^{FLAG}$DNABJ8$^{H31Q}$ and the other plate was transfected with 5 µg of DNA encoding GFP. Both plates were plates were split into 6-well plates or three separate 6 cm plates that were previously coated with Poly-D-Lysine. Each individual well was incubated with an increasing concentration (0 µM, 25 µM, 50 µM, 100 µM, 200 µM, 500 µM) of a metal for 15 min at 37 °C. Toxic metal media was replaced with fresh media and plates were incubated for 16 h at 37 °C to recover. Plates were then harvested by scraping in DPBS and then lysed in 9:1 RIPA:10x PIC in ice for 30 min.

Lysates were separated from cell debris by centrifugation at 21,000 *x g* for 15 minutes at 4 °C. Bradford assay was used to quantify protein concentration in each lysate. 2 µg/µl of protein samples were prepared for western blot analysis after addition of 9:1 Laemlli:1 M DTT (17% of solution).

15 µg of each samples were separated on a 12% SDS-PAGE Gel with 1.0 mm thickness. Western blots were probed for Flag (Sigma M2 monoclonal anti-Flag antibody), HSPA1A (rabbit polyclonal), Beta-actin (mouse monoclonal 7D2C10), and GFP (rabbit polyclonal) primary antibodies, near-IR secondary antibodies (Li-COR) and visualized on a Li-Cor Biosciences Fc Imager.

<u>Assessing E1-E2 interaction in PDC Complex after Arsenite Treatment</u>

*Immunoprecipitation of $^{Flag}$PDHA1*: For each two plex, four 10 cm plates of HEK293T cells were transfected by the calcium phosphate method with 5 µg of plasmid DNA encoding $^{Flag}$PDHA1 in the pFLAG backbone[1]. Briefly, 5 µg of DNA in 1 mL 250 µM $CaCl_2$ is vortexed while adding dropwise 1 mL HBS 2X for 10 seconds at ambient temperature, the transfection solution is promptly (≤ 15 min) added dropwise to cells, and the cell media is changed between 12 and 16 h. A positive transfection control is performed with GFP alongside each transfection. Plates were treated with heavy metal salts or vehicle at 40-46 hours post transfection. Cells were harvested by scraping in DPBS. Cells were diluted to 1ml of DPBS and incubated and rotated with 1 mM DSP (dithiobis(succinimyidyl propionate) in 1% DMSO vehicle for 30 mins at ambient temperature. DSP was quenched with 100 mM Tris pH 8 (final concentration) and rotated for 15 minutes at room temperature. Cells were then lysed in 9 parts RIPA Buffer (150 mM NaCl, 50 mM Tris pH 7.5, 1% Triton X-100, 0.5% sodium deoxycholate, 0.1% SDS) and 1 part 10x PIC for 30 min on ice. Lysate was separated from cell debris by centrifugation at 21,000 *x g* for 15 min at 4 °C. Protein in the lysate was quantified by Bradford. Each sample had protein content normalized to same amount and diluted to 3 µg/µl. Lysates were pre-cleared with 15 µL Sepharose-4B beads for 30 min at 4 °C, then centrifuged at 1,500 *x g* for 1 min to pellet beads. Lysate was then separated and incubated with 15 μL of M2 anti-Flag Magnetic Beads and rotated overnight at 4 °C. The anti-Flag beads were washed the next day four times with RIPA buffer. Each wash included rotation for 10 minutes at ambient temperature. Proteins bound to the anti-Flag beads were eluted by boiling for 5 min at 100 °C in 25 μL of Laemmli concentrate (120 mM Tris pH 6.8, 60% glycerol, 12% SDS, brilliant phenol blue to color). Eluates were blotted for Western Blot on 10% SDS-PAGE Gel with 1.0 mm thickness. Western blots were first probed to observe E1-E2 interaction: 1. Rabbit polyclonal anti-DLAT, 2. Rabbit polyclonal anti-PDHB, 3. Rabbit polyclonal anti-PDHA1. Blots were then probed with mouse monoclonal M2 anti-Flag (Sigma), mouse monoclonal anti-Beta Actin (7D2C10), and rabbit polyclonal anti-GFP using near-IR secondary antibodies (Li-COR). Bands corresponded to each antibody were visualized and quantified by densitometry on a Li-Cor Biosciences Fc Imager using Image Studio. DLAT and PDHB band intensities were normalized to band intensities from the bait, PDHA1.

### SUPPLEMENTAL RESULTS AND DISCUSSION

Robustness of Hsp40 affinity profiling by AP-MS:

The Hsp40 Affinity mass spectrometry experiments are comparisons between two sets of twelve AP-MS preparations, analyzed as four individual multiplexed injections of six samples each. To indicate the robustness of hits from this assay, the Strictly Standardized Mean Differences (SSMD) were compared between each multiplexed injection for both arsenic (**Figure S7**) and cadmium **(Figure S8**). SSMDs represent the mean differences between the treated and untreated populations, normalized by root-mean-square standard deviations. The most affected proteins reproduce well across each replicate. It is worth noting that neither arsenite nor cadmium treatment affects the bulk proteome of eluted DNAJB8$^{H31Q}$ co-IPs by silver stain of the eluate (**Figures S9** and **S10**).

Comparison with DNAJB8 interactor studies

The arsenite pulldowns collectively identified and quantified 1696 unique proteins, not including 28 keratin/immunoglobulin proteins that were excluded from the set, and 24 proteins that were identified but whose TMT reporter ions could not be quantified. We previously characterized 562 high confidence DNAJB8$^{H31Q}$ interactors from AP-MS in unstressed HEK293T cells[1]. Those included 379 of the proteins found in this study, and 21/30 of our high confidence arsenite-sensitive proteins. It is possible that stress conditions, by increasing the affinity and thus recovery of a select proteome, also increase the likelihood of identifying those proteins from data-dependent analysis.

Piette et al recently[18] reported comprehensive interactomes of human Hsp40 and Hsp70 family proteins (unlike our work, these all had active J-domains), finding 37 high confidence interactors of DNAJB8 and another 479 proteins that could not be reliably distinguished from background. We find 22/37 of their high-confidence interactors in our data set, with most of the missing proteins being either Hsp70s (which bind through the J-domain) or Hsp70 binding proteins (Hsp40s, CHIP, FKBP8 etc.). Only

11/31 of our high confidence arsenite-sensitive proteins are found in their study, and none are among their high confidence interactors.

Optimization of LiP assay for As target proteins

To compare proteolytic susceptibilities over a wide dynamic range, we incubated lysates in varying amounts of PK. Proteins that are destabilized due to cellular arsenite exposure should have regions that are more solvent exposed, and hence more sensitive to PK digestion compared to untreated samples. PK resistance in proteins has been used to measure stability of native proteins after exposing them to a PK gradient and thus PK concentrations can be used as a scale to measure changes in a protein's structure[19].

The PK concentration gradient was optimized to bracket the full range of changes in observable protein on a Coomassie-stained SDS-PAGE gel (**Figure S11**). Protein band intensity decreases around 1:1000 PK:protein ratio and little intact protein is observable at 1:100 PK:protein. These values are somewhat lower than commonly reported conditions, which range from as 1:33 to 1:100[20,21]. Peptides for monitoring were chosen in the Picky software, based on chemical properties amenable to PRM and to maximize distribution across the chromatographic gradient.

We determined assay precision across technical and biological replicates. To determine the reproducibility of the scheduled PRM assay itself, ten consecutive injections of a lysate tryptic digest were performed, and CVs calculated for the integrated fragment intensities of the 13 targeted peptides from a trypsin only digested lysate. Each targeted run included a 6 µg injection, a 10 millisecond retention time window for each peptide, and chromatograms that were scanned at 7500 nominal resolving power. The low resolving power was chosen to minimize cycle time. Lower cycles times allow more scans across peaks, increasing robustness of quantification[22]. The MS2 intensities of transitions from a precursor ion were integrated by Skyline for quantification of each peptide[21]. The median CV for the summed transitions MS2 among the 13 targeted peptides was 20%. The median CV for the summed MS2 transitions normalized to internal standard was only slightly improved to 14% (**Supp. Table 4**). These CVs are typical for PRM[23,20]. TDYNASVSVPDSSGPER (HNRNPK) had a CV greater than 30%, possibly due to their hydrophilic character as they eluted early from the reversed-phase $C_{18}$ column[24].

To determine whether 7500 nominal resolving power was too low to avoid matrix interference[17], we performed 6 more technical replicate PRM experiments on a HEK239T tryptic digest using 60,000 nominal resolving power and scheduling only three select peptides and the internal standard. The four peptides selected had 10 minute retention time windows that did not overlap with each other to minimize cycle times. The calculated CV from the six runs were compared with the quantification of the previous measurements that were run at 7500 resolution. The median CV for the summed transitions MS2 among the 4 targeted peptides was 18. The median CV for the summed MS2 transitions normalized to internal standard among the 4 peptides was 9% (**Supp. Table 5**). Higher resolving power hence did not meaningfully increase the precision of the method, consistent with a recent report on the dispensability of accurate mass for PRM[20].

The CV between biological replicates was addressed during the LiP mass spec runs. The LiP mass spec samples included three zero PK controls and the following Protein: PK samples: 5000:1, 2000:1, 1000:1, 500:1, 200:1, and 100:1. Peptides were scanned in 10 minute retention time windows at 7500 resolution after a 6 ug HEK293T tryptic digest injection. The CV was calculated among the three Zero PK samples for both arsenite exposed and control cellular conditions. Across these biological replicates, median CV values were below 20% with only 1 peptide having a CV greater than 25% (**Supp. Table 6**). The median CV for LiP method targeting TDP-43 peptides was below 17%. (**Supp. Table 7**). The E1 subunit LiP experiment that looked at 11 peptides between PDHA1 and PDHB found CV below 20% for all peptides (**Supp. Table 8**). In summary, the peptide CVs in our PRM assay for biological replicates were relatively similar to the CVs calculated for technical replicates.

Investigating the Interaction of E1 and E2 subunit in Pyruvate Dehydrogenase Complex

Both AP-MS and LiP experiments reveal that PDHA1 and PDHB proteins misfold after arsenite treatment. Both PDHA1 (fold change = 1.5 ± 0.4, q = 2 x $10^{-4}$) and PDHB (fold change = 1.3 ± 0.3, q = 0.06) increase their affinity to $^{FLAG}$DNAJB8$^{H31Q}$ after cellular treatment with 500 µM sodium meta arsenite ($NaAsO_2$) for 15 min. Seven peptides in the PDHA1 and PDHB proteins have increased susceptibility to proteinase K after this arsenite treatment (**Figure S3C**).

The evidence of protein misfolding for both these proteins by AP-MS and LiP prompted further investigation of understanding the effects of arsenite on the Pyruvate Dehydrogenase Complex (PDC), particularly understanding the role of a destabilized E1 subunit. PDC consists of an icosahedral core of E2 subunits peripherally decorated with E1 and E3 subunits. Two PDHB and two PDHA1 proteins comprise each heterotetrameric E1 subunit, which decarboxylates pyruvate to acetylate a lipoamide cofactor[25]. The lipoamide cofactor is covalently bound to the core E2 subunit, which consists of several dozen copies of DLAT along with regulatory proteins[26]. DLAT catalyzes transfer of the acetyl group from lipoamide to synthesize acetyl-CoA, the primary feedstock for cellular respiration. Destabilization of E1 activity is consistent with impaired PDC activity and consequently impaired mitochondrial ATP production, a major consequence of cellular arsenic exposure[27]. Direct arsenite binding to lipoamide could affect E1 stability if it impaired lipoamide interactions with the E1 active site[28]. However, we did not see any change in PK susceptibility near the lipoamide binding site of E1 (**Figure S4**). An alternative mechanism for E1 misfolding is that arsenite treatment affects the salt bridges that bind E1 to the peripheral subunit binding domain (PSBD) of E2[29]. The IMEGPAFNFLDAPAVR peptide located at the C-terminus of PDHB, which participates in PSBD binding, is more sensitive to Proteinase K after arsenite treatment (**Figure S4**). Loss of interaction at the PSBD domain could lead to DLAT dissociating from the E1 subunit as a result of arsenite treatment, the PDC.

We evaluated PDC stability by co-IP of E2 with E1[30]. $^{Flag}$PDHA1 was overexpressed in HEK293T cells, which were then treated with either arsenite (500 μM for 15 min) or with water (control), lysed, and immunoprecipitated with anti-Flag beads. Eluates were analyzed by SDS-PAGE and immunoblotting. Crosslinking with 1 mM DSP crosslinking following cell harvest provided optimal recovery of PDHB and DLAT (**Figure S12**). No significant differences in DLAT binding to E1 were observed (**Figures S5 and S13**). Literature reports achieve stoichiometric inhibition of purified pyruvate dehydrogenase with lower concentrations of trivalent arsenite and longer times[31]. We treated HEK293T cells overexpressing $^{Flag}$PDHA1 with 2 μM arsenic for four hours. No discernable differences were seen in DLAT binding (**Figures S6 and S14**). Hence, even though the E1 complex is destabilized by arsenite exposure, the destabilization does not induce complex disassembly.

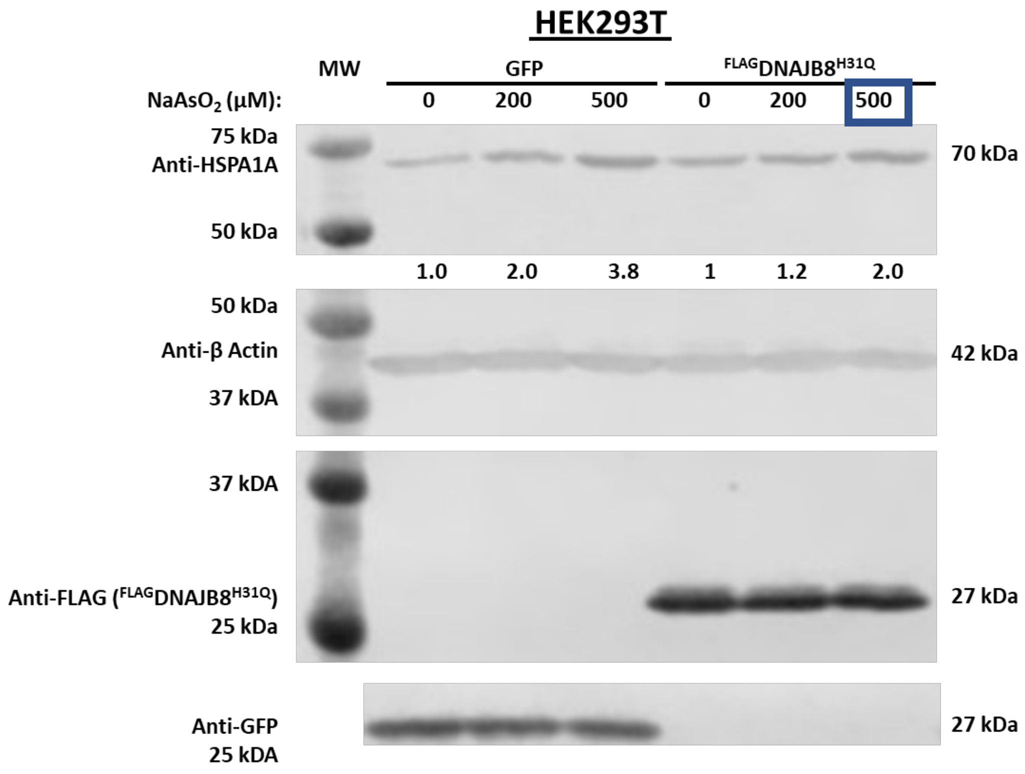


**Figure S1:** Western Blot analysis of HSR induction by arsenite on HEK293T Cells. Cells were treated for 15 min. with the indicated concentration of $NaAsO_2$ and then allowed to recover for 16 h to allow for accumulation of stress-induced proteins. 500 µM $NaAsO_2$ was chosen for AP-MS experiments and limited proteolysis experiments. Predicted weights for antibodies are shown on the right. Numerical values below Anti-HSPA1A slice are band intensities normalized to the 0 µM condition. Antibody for GFP is shown on 800 channel only.

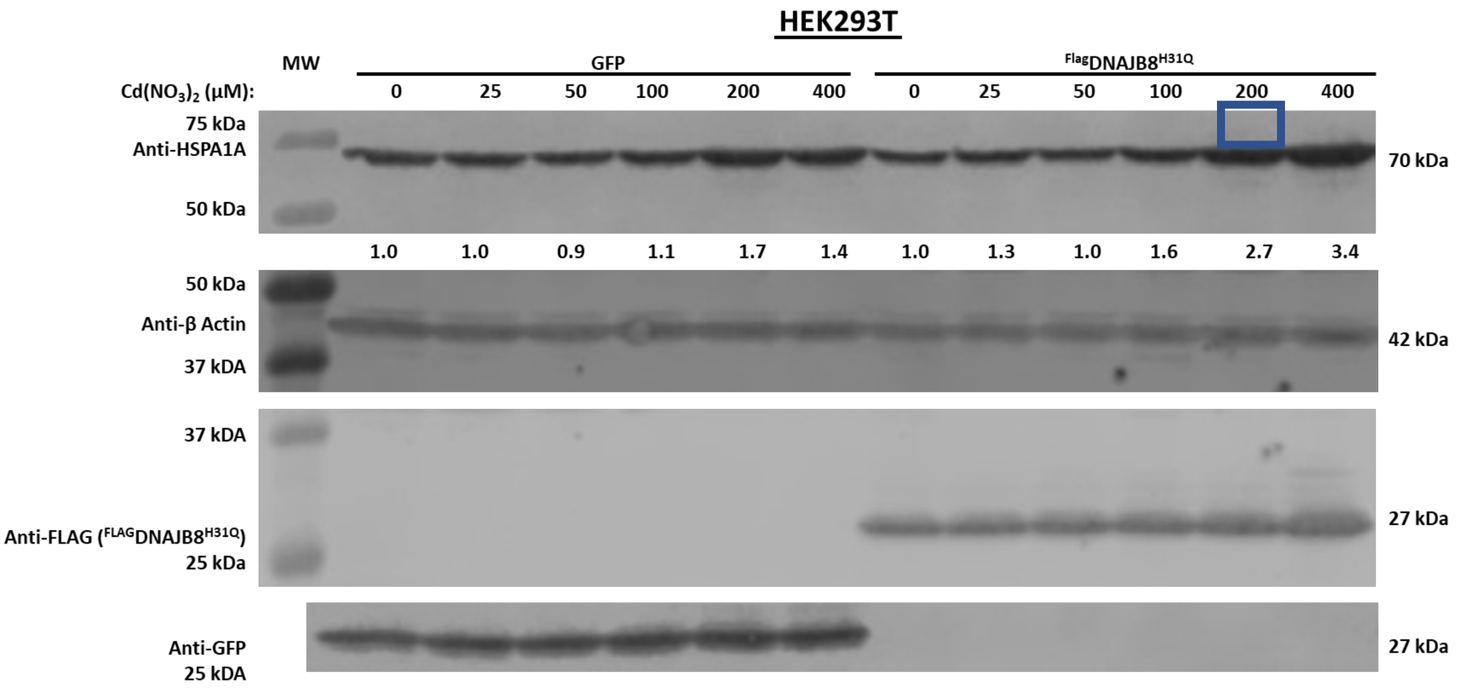


**Figure S2:** Western Blot analysis of HSR induction by cadmium on HEK293T Cells. Cells were treated for 15 min. with the indicated concentration of $Cd(NO_3)_2$ and then allowed to recover for 16 h to allow for accumulation of stress-induced proteins. 200 µM $Cd(NO_3)_2$ was chosen for AP-MS Experiments. Predicted weights for antibodies are shown on the right. Numerical values below Anti-HSPA1A slice are band intensities normalized to the 0 µM condition. Antibody for GFP is shown on 800 channel only.

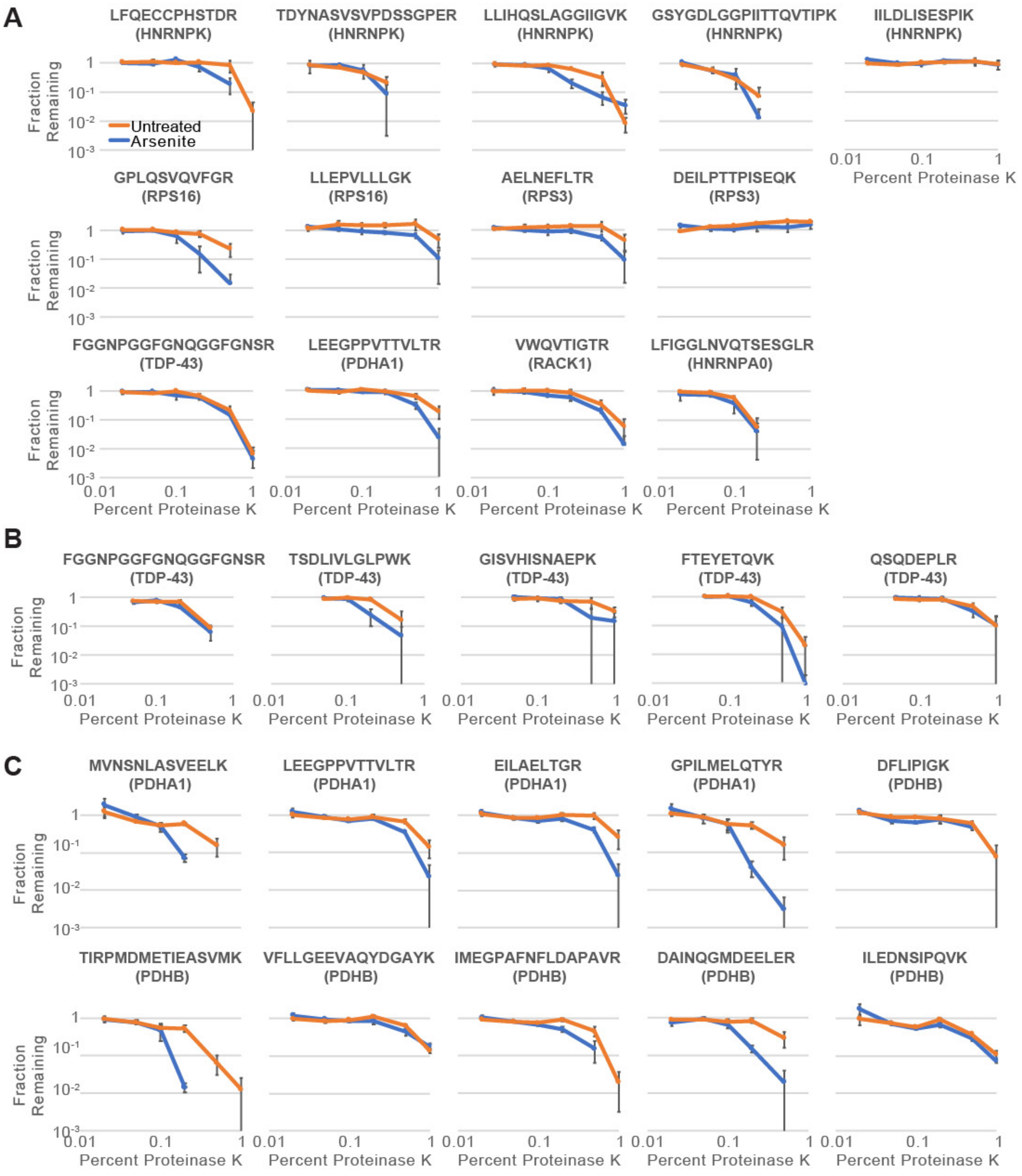


**Figure S3**: **A**) Proteinase K susceptibility curves for 13 peptides for several protein targets from the Hsp40 affinity assay. **B**) Proteinase K susceptibility curves for TDP-43 peptides. **C**) Proteinase K susceptibility curves for PDHA1 and PDHB peptides. Samples from untreated cells are in orange and samples from arsenite-treated cells are in blue (500 µM, 15 min). (n = 3 biological replicates). LC-PRM runs were performed in technical triplicate for set **A,B** and averaged. Error bars represents standard error across biological replicates. Representative chromatographic traces from Skyline are presented in **Figure S15**.

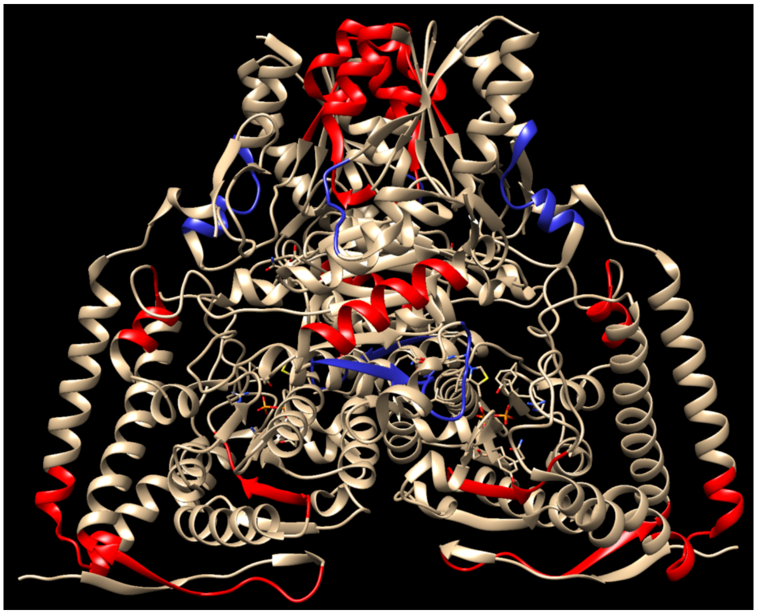

**Figure S4**: Location of proteinase K sensitive peptides after arsenite treatment are shown in the E1 subunit (PDB entry, 1N14). Peptides more sensitive to proteinase K after treatment are shown in red and peptides that did not change in sensitivity after treatment are in blue.

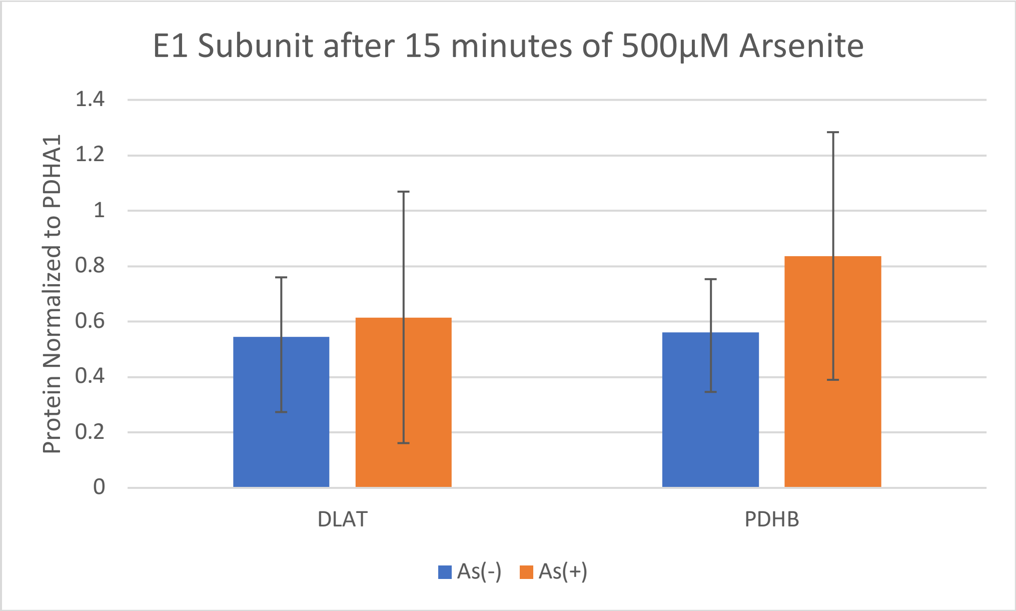


**Figure S5.** 15 minutes of 500 µM $NaAsO_2$ did not show evidence of E1 dissociating from E2 subunit. HEK293T cells transfected with $^{Flag}$PDHA1 were treated with 500 µM of arsenite for 15 minutes prior to lysis. Protein eluates obtained after immunoprecipitation of $^{Flag}$PDHA1 were blotted on SDS-Page gels. Band intensities of PDHB and DLAT were normalized to $^{Flag}$PDHA1 (bait). Average of normalized PDHB and DLAT among all four replicates are shown above. No significant changes in amount of DLAT binding and PDHB binding to $^{Flag}$PDHA1 after arsenite treatment was observed (from n = 4 biological replicates). Error bars represent standard deviation. A representative Western blot is shown in **Figure S11**.

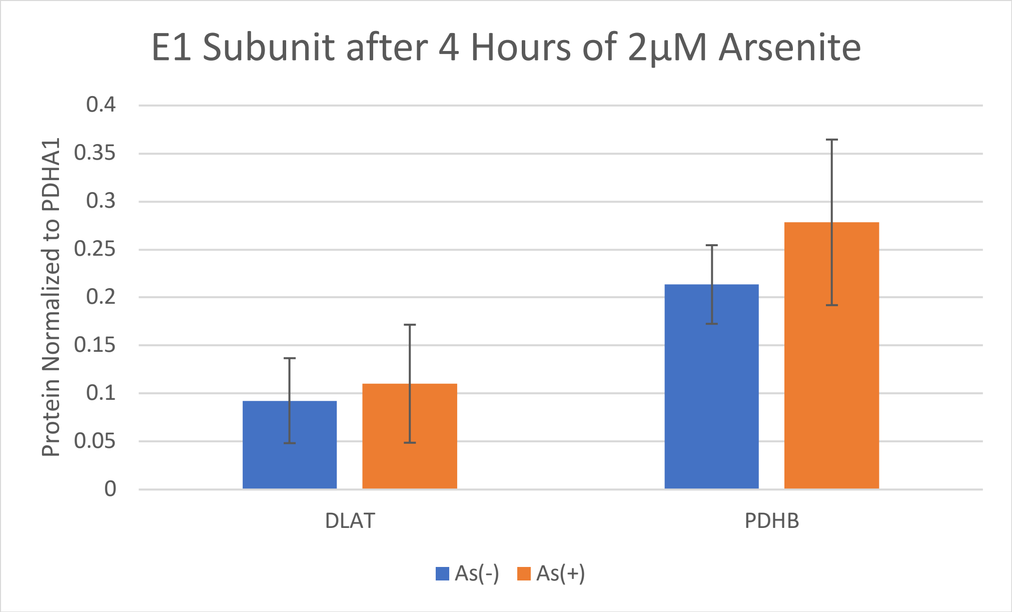


**Figure S6.** 4 hours of 2 µM $NaAsO_2$ did not lead to DLAT dissociating from E2 subunit. HEK293T cells transfected with $^{Flag}$PDHA1 were treated with 2µM of arsenite for 4 hours prior to lysis. Protein eluates obtained after immunoprecipitation of $^{Flag}$PDHA1 were blotted on SDS-Page gels. Band intensities of PDHB and DLAT were normalized to $^{Flag}$PDHA1 (bait). Average of normalized PDHB and DLAT among all four replicates are shown above. No significant change in amount of DLAT binding and PDHB binding to $^{Flag}$PDHA1 after arsenite treatment was observed (from n = 4 biological replicates). A representative Western blot is shown in **Figure S12.**

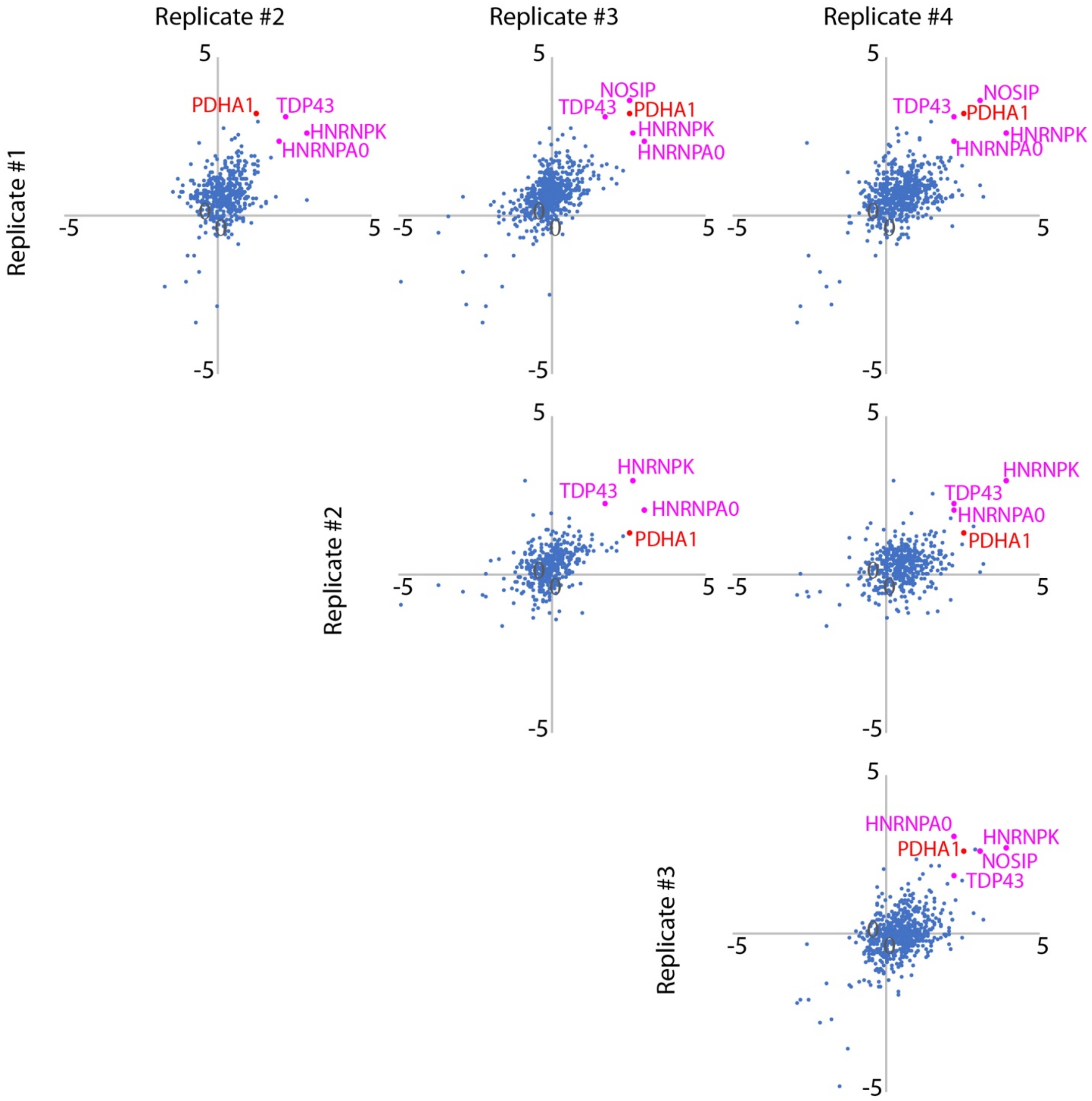


**Figure S7**: Strictly Standardized Mean Differences of individual TMT-AP-MS experiments involving cellular treatment by arsenite (500 µM, 15 minutes). Each plot compares separate runs (each run includes three arsenite-treated biological replicates and three vehicle-treated biological replicates) and includes fold changes that reflect the ratios between DNAJB8-normalized integrated TMT reporter ion intensities for the control (vehicle) and arsenite-treated cells. TDP43 and PDHA1 are labeled as pink and red and consistently increase in affinity to DNAJB8$^{H31Q}$ after arsenite treatment.

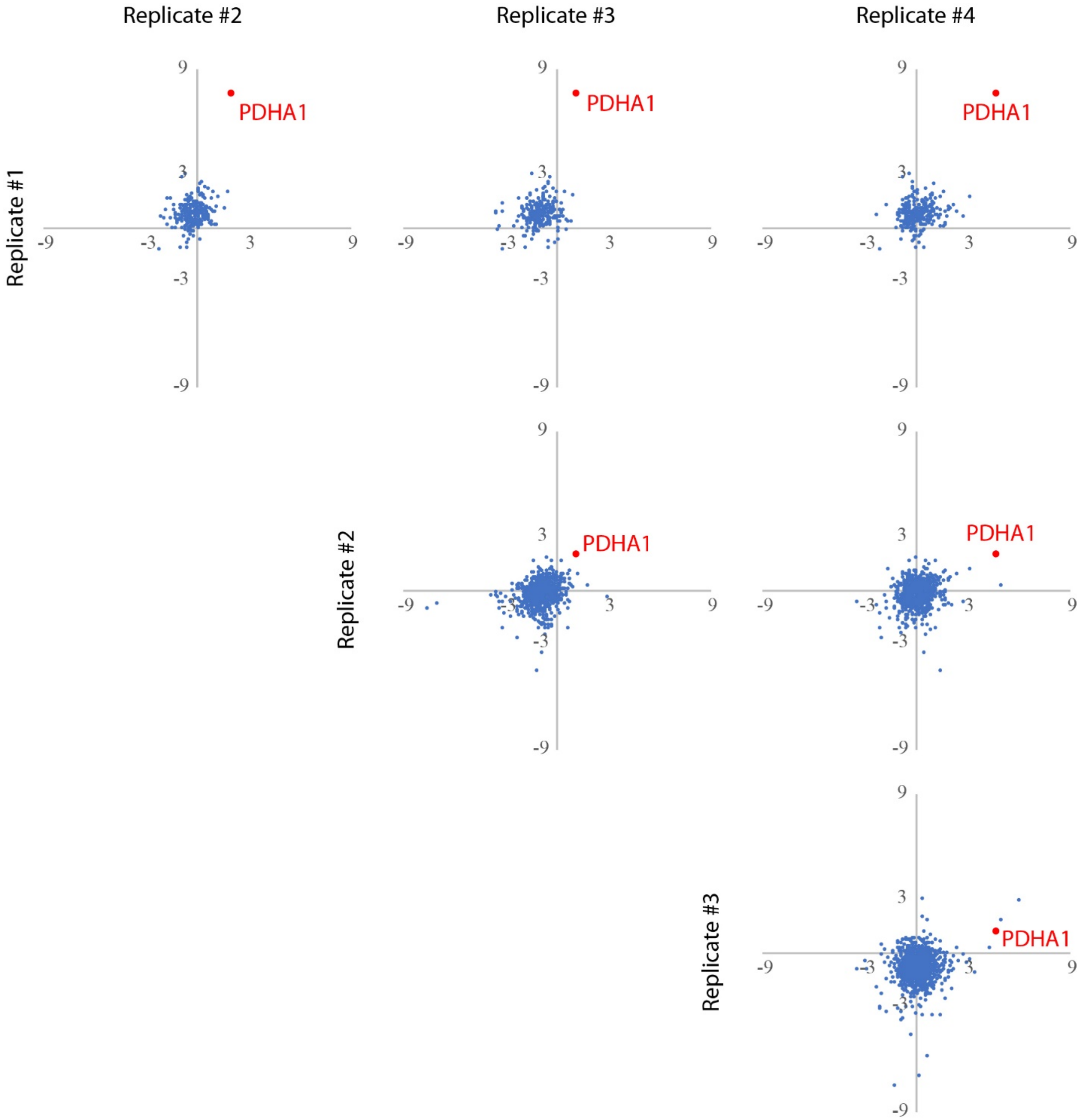


**Figure S8**: Strictly Standardized Mean Differences of individual TMT-AP-MS experiments involving cellular treatment by cadmium (200 µM, 15 minutes). Each plot compares separate runs (each run includes three cadmium-treated biological replicates and three vehicle-treated biological replicates) and includes fold changes that reflect the ratios between DNAJB8-normalized integrated TMT reporter ion intensities for the control (vehicle) and cadmium-treated cells. PDHA1 is labeled as red and consistently increase in affinity to DNAJB8$^{H31Q}$ after cadmium treatment.

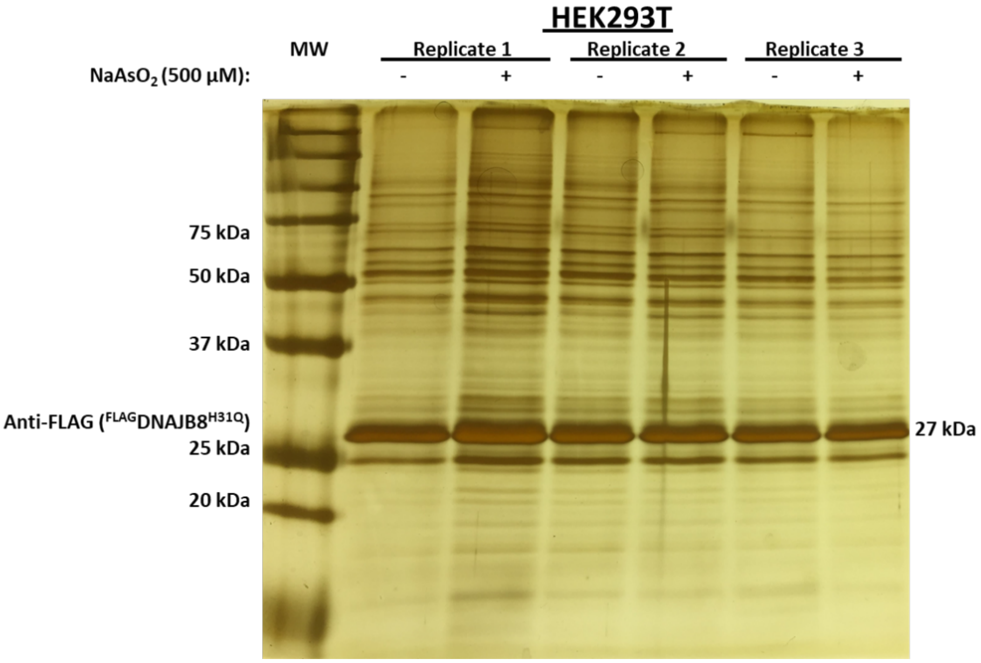


**Figure S9:** Representative Silver Stain for an Arsenite AP-MS. Each replicate contained two transfected ${}^{FLAG}$DNAJB8${}^{H31Q}$ 10cm plates treated with either 500 µM $NaAsO_2$ or water (control). Three replicates are stained to show bait and visually show differences in prey after each respective pulldown. ${}^{Flag}$DNAJB8${}^{H31Q}$ is the most abundant protein in each replicate after the immunoprecipitation. Other bands represent proteins recovered with DNAJB8.

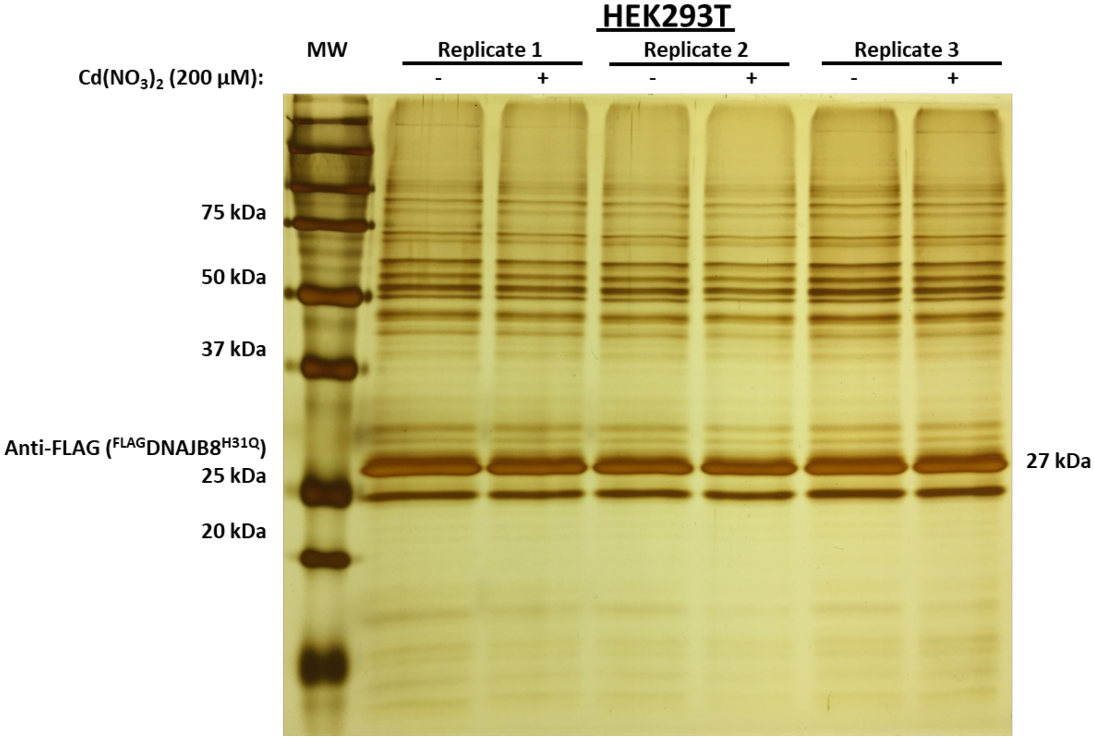


**Figure S10**: Representative Silver Stain for a Cadmium AP-MS. Each replicate contained two transfected $^{FLAG}$DNAJB8$^{H31Q}$ 10cm plates treated with either 200 µM $Cd(NO_3)_2$ or water (control). Three replicates are stained to show bait and visually show differences in prey after each respective pulldown. $^{Flag}$DNAJB8$^{H31Q}$ is the most abundant protein in each replicate after the immunoprecipitation. Other bands represent proteins recovered with DNAJB8.

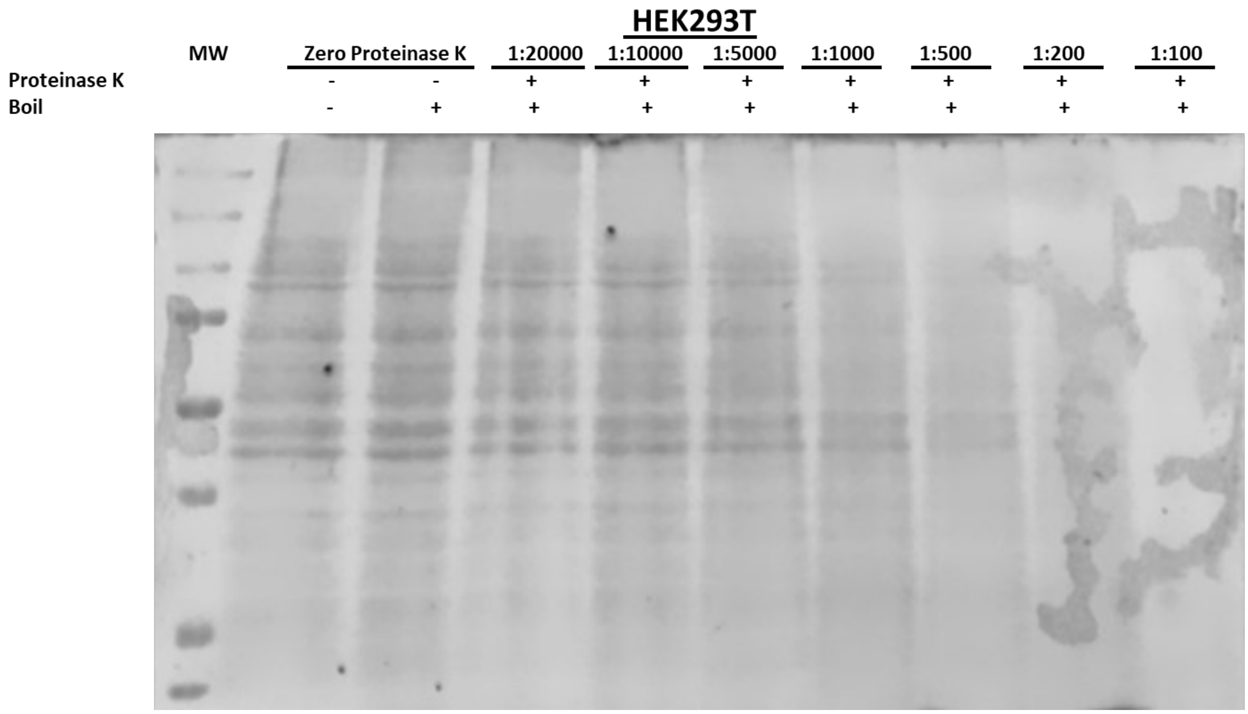


**Figure S11**: Coomassie Stain of Proteinase K Optimization. Each lane represents a different Proteinase K to Protein ratios. Samples of lysates were incubated with an amount of proteinase K for 1 minute at 25 degrees and then boiled for 5 minutes. Protein bands are no longer visible in the 1:100 Proteinase K: Protein sample. 25ug of each lysate was used.

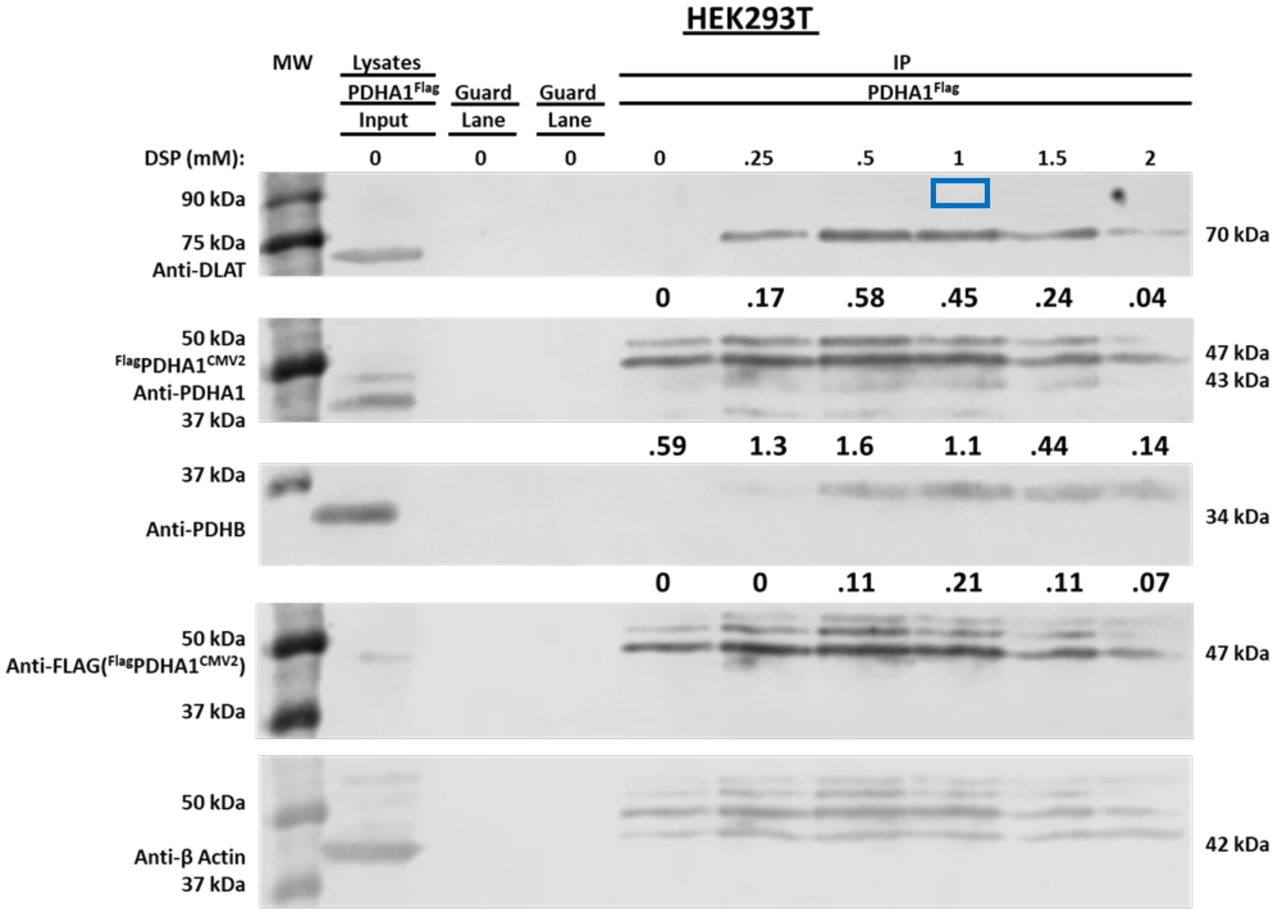


**Figure S12**: Western Blot Analysis for effects of DSP crosslinking on pulldown of E1 and E2 subunit. 1mM of DSP crosslinker was chosen as conditions for the immunoprecipitation of $^{Flag}$PDHA1. Transfected HEK293T cells were incubated with DSP crosslinking during cell harvest. Predicted weights are shown on the right. Numerical values shown below Anti-DLAT slice, Anti-PDHA1 and Anti-PDHB are densitometric intensities determined using Li-COR Biosciences Image Studio.

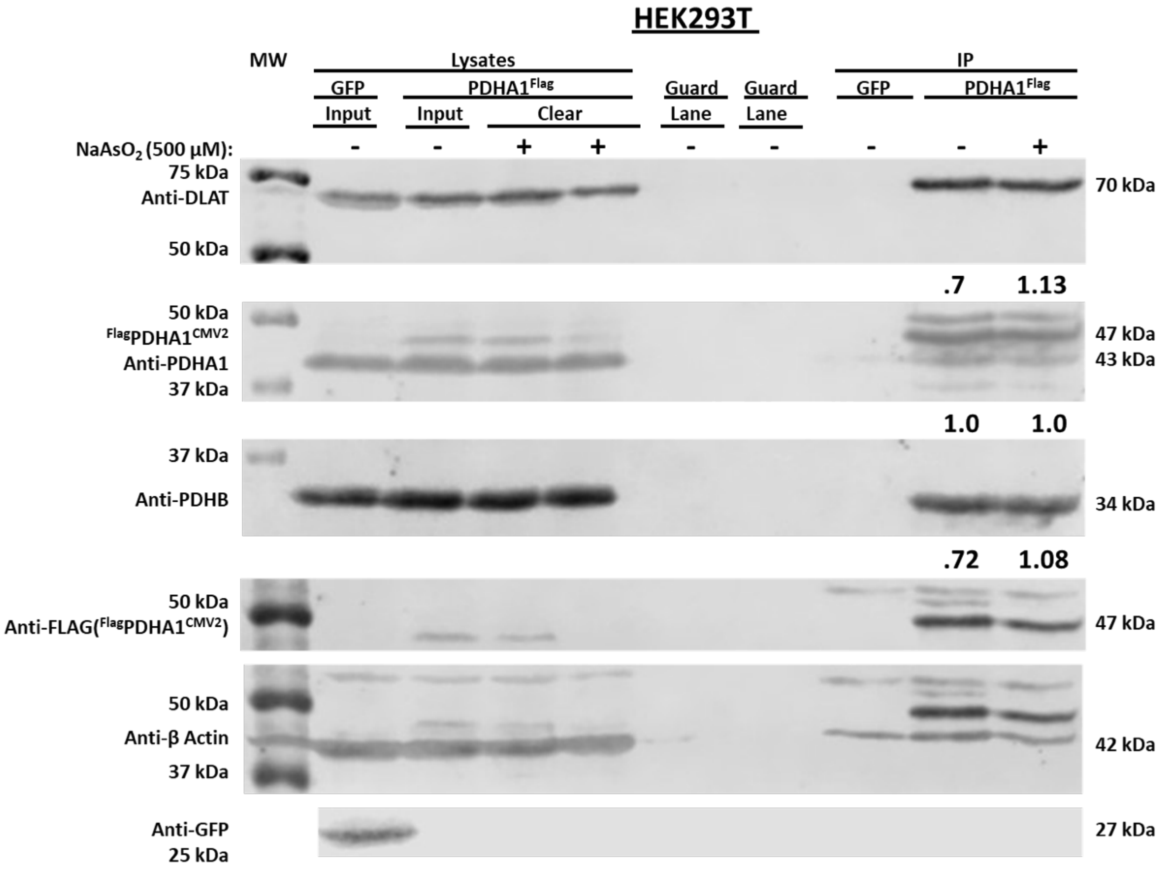


**Figure S13**: Representative western blot analysis for the effects of Arsenite on E1 and E2 subunit interaction after 15 minutes of treatment. One of four biological replicates is shown above. Transfected HEK293T cells were incubated with 500 µM $NaAsO_2$ for 15 minutes at 37 $^oC$. Predicted weights for antibodies are shown on the right. Numerical values below Anti-DLAT, Anti-PDHA1, and Anti-PDHB slice are band intensities normalized to the intensity of PDHA1. Antibody for GFP is shown on 800 channel only.

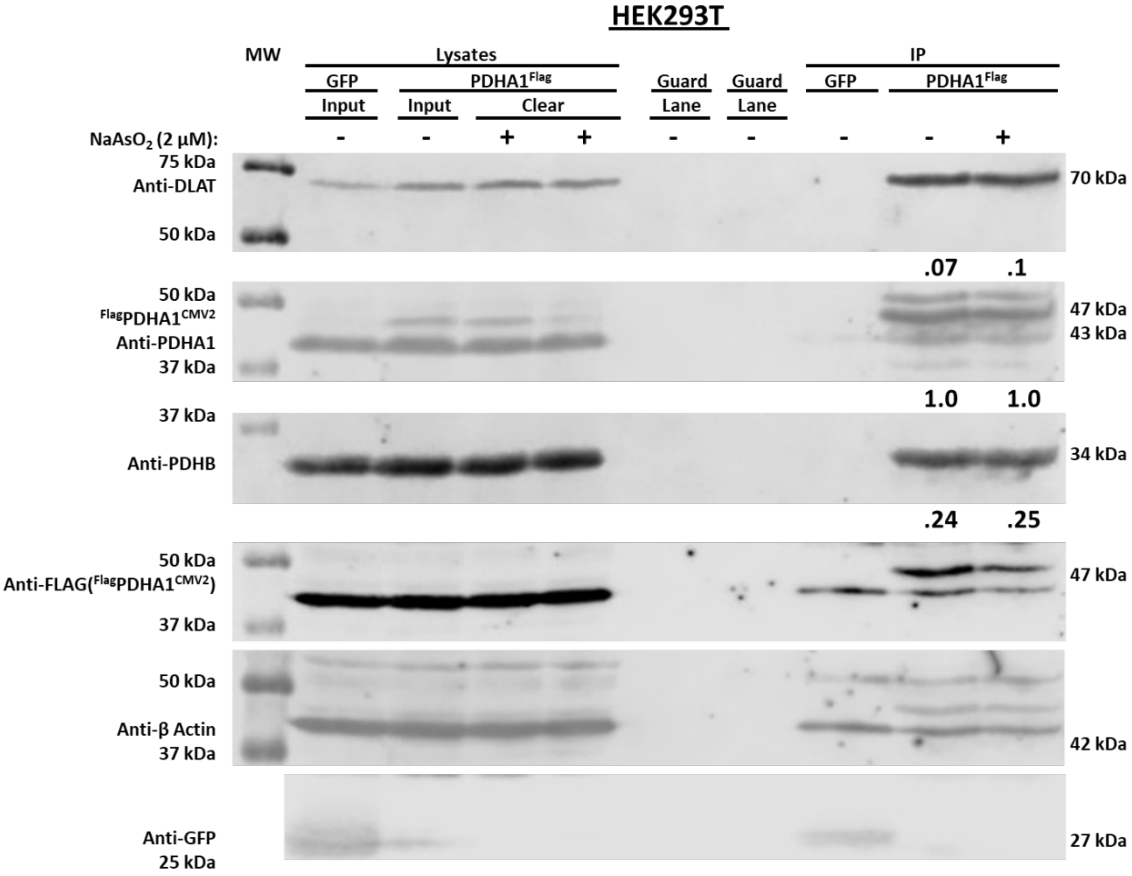


**Figure S14**: Representative Western Blot analysis for Arsenite effects on E1 and E2 subunit interaction after 4 hours of treatment. One of four replicates is shown above. Transfected HEK293T cells were incubated with 2 µM $NaAsO_2$ for 4 hours at 37 $^oC$. Predicted weights for antibodies are shown on the right. Numerical values below Anti-DLAT, Anti-PDHA1, and Anti-PDHB slice are band intensities normalized to the intensity of PDHA1. Antibody for GFP is shown on 800 channel only.

**Fig. S15.** Representative Chromatograms of selected peptides for LiP.

RPS3, DEILPTTPISEQK, 735.888 2+:

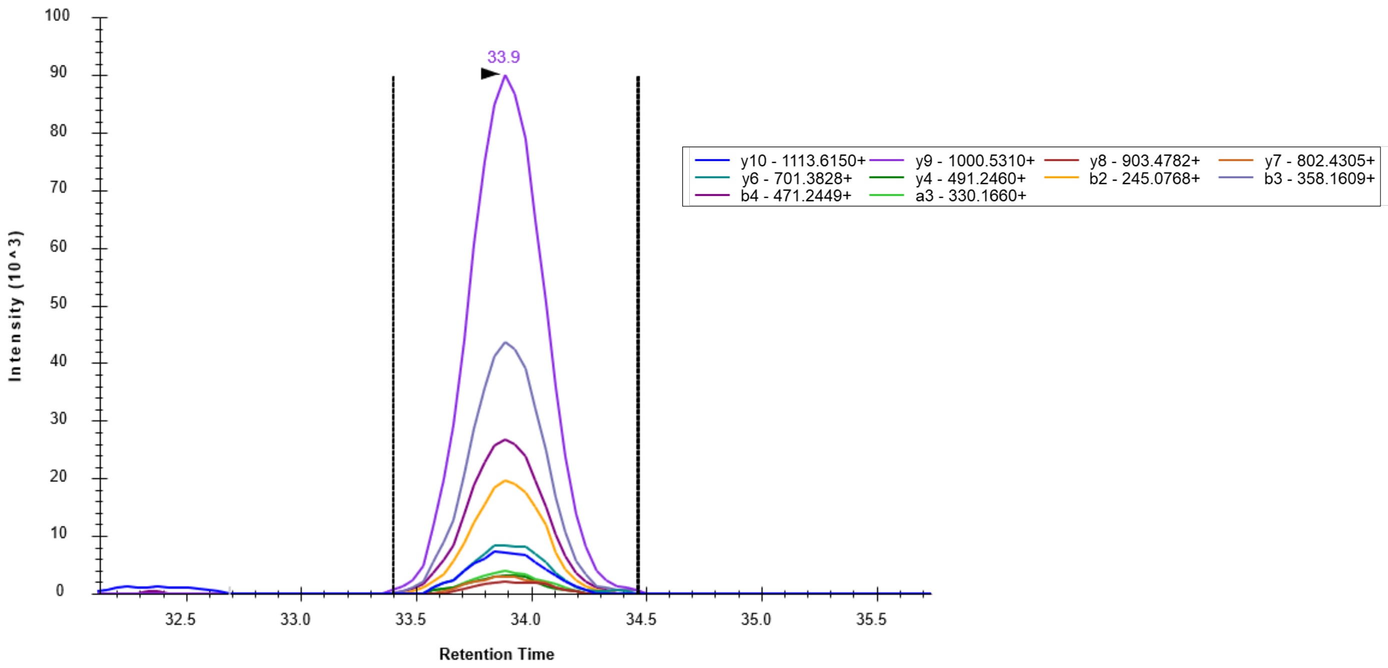


RPS3, AELNEFLTR, 546.78784 2+:

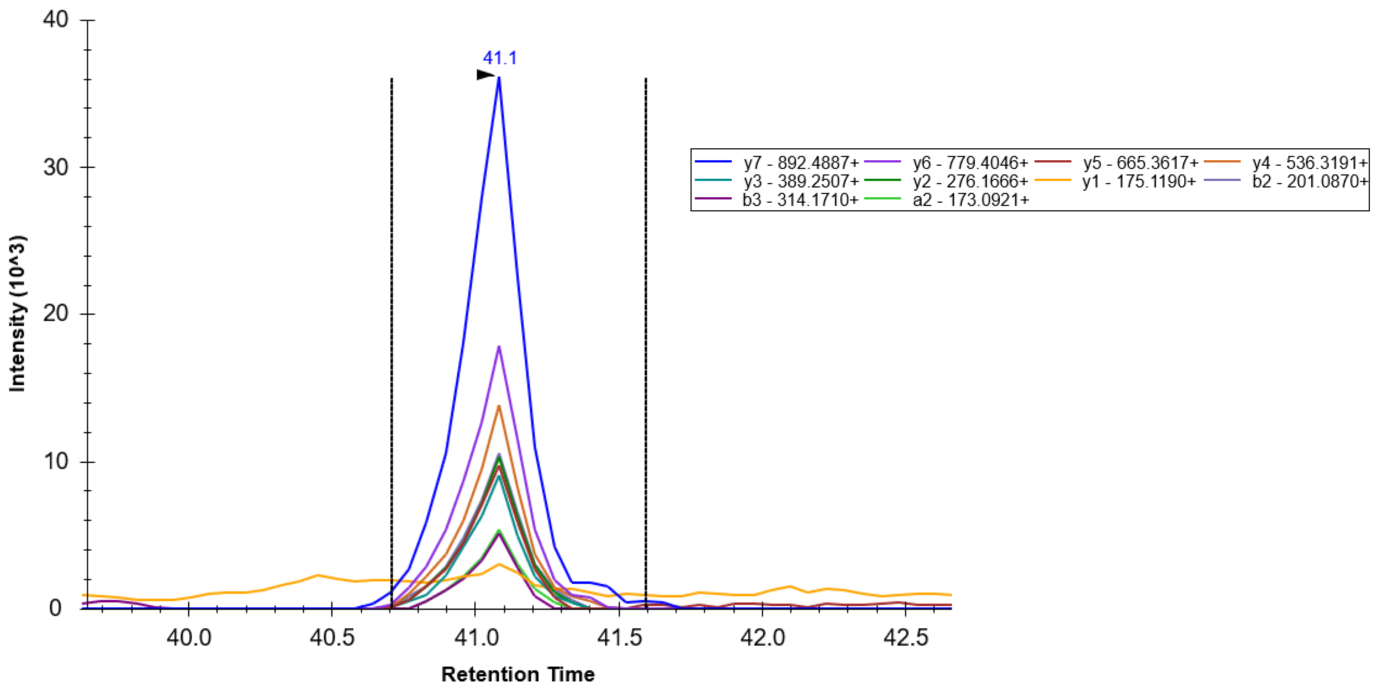

HNRNPK, IILDLISESPIK, 670.90541 2+:

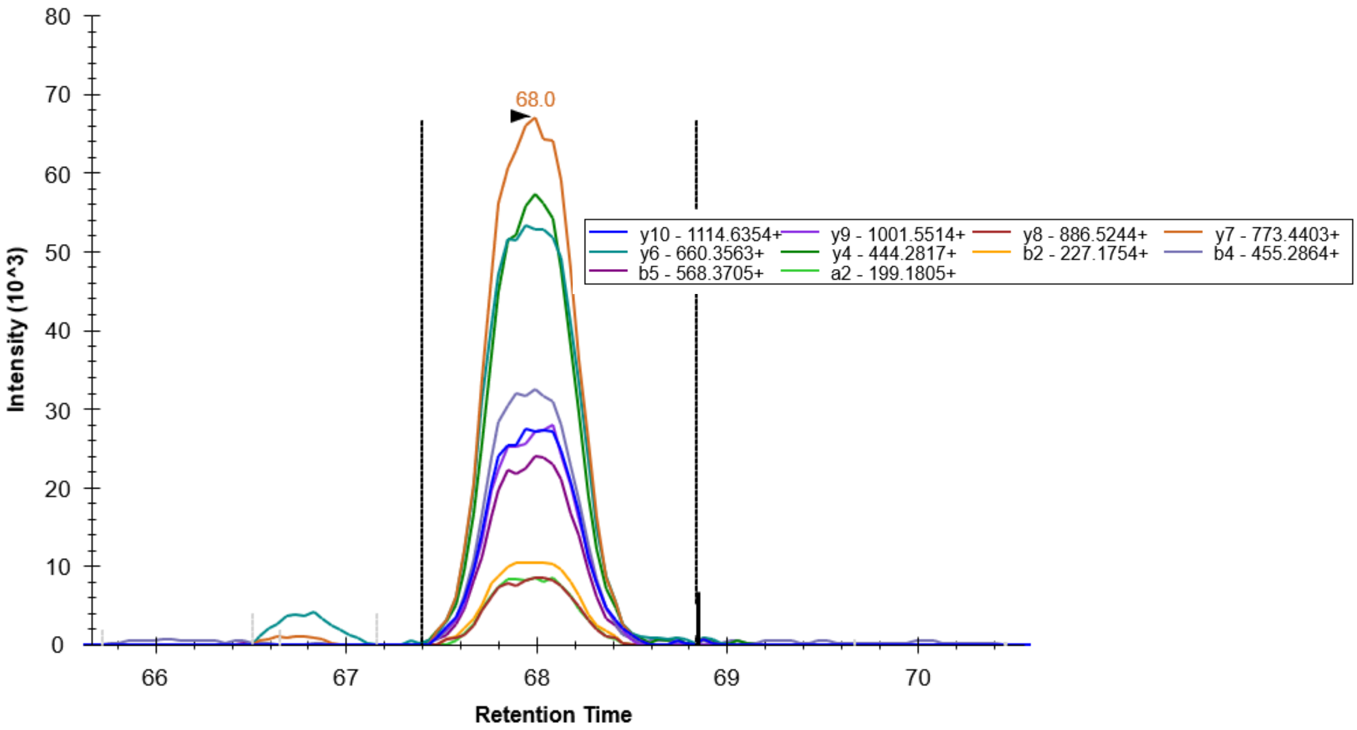


HNRNPK, GSYGDLGGPIITTQVTIPK, 959.02002 2+:

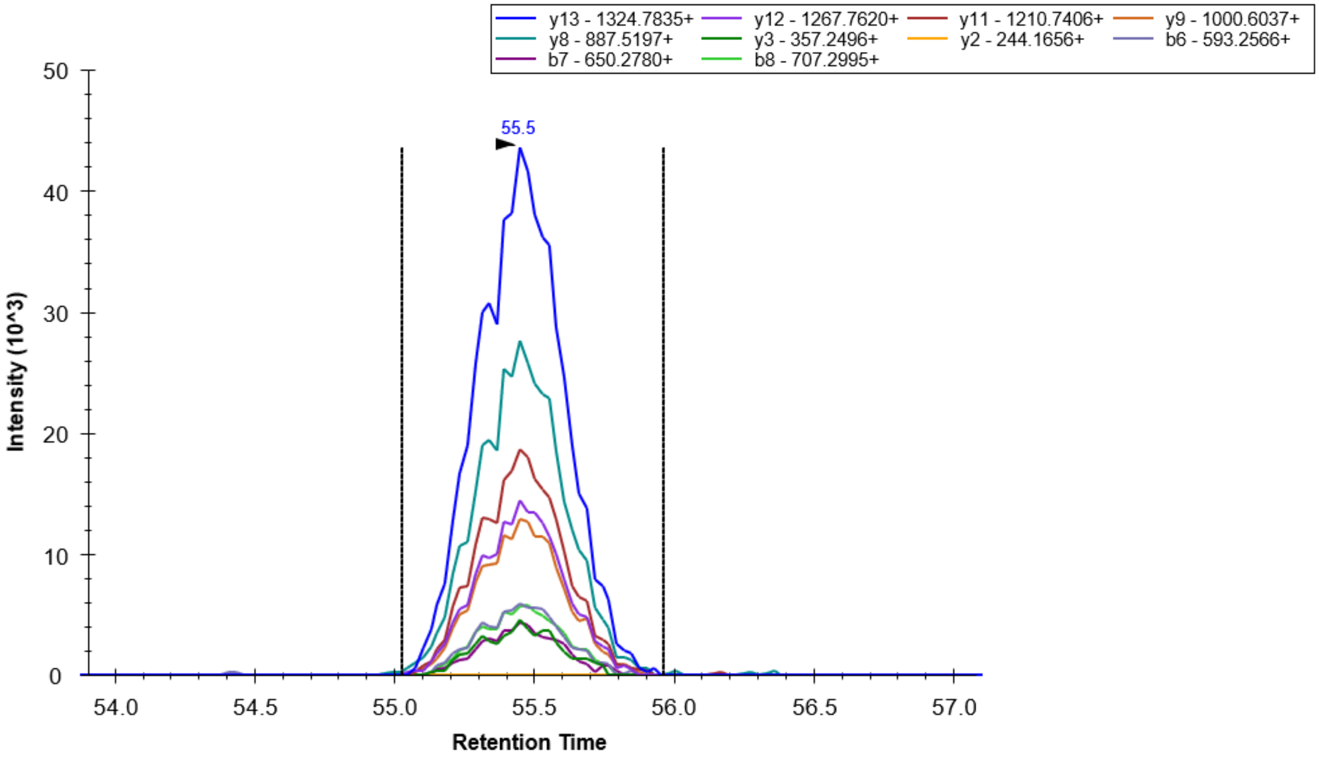

HNRNPK, LLIHQSLAGGIIGVK, 759.97195, 2+:

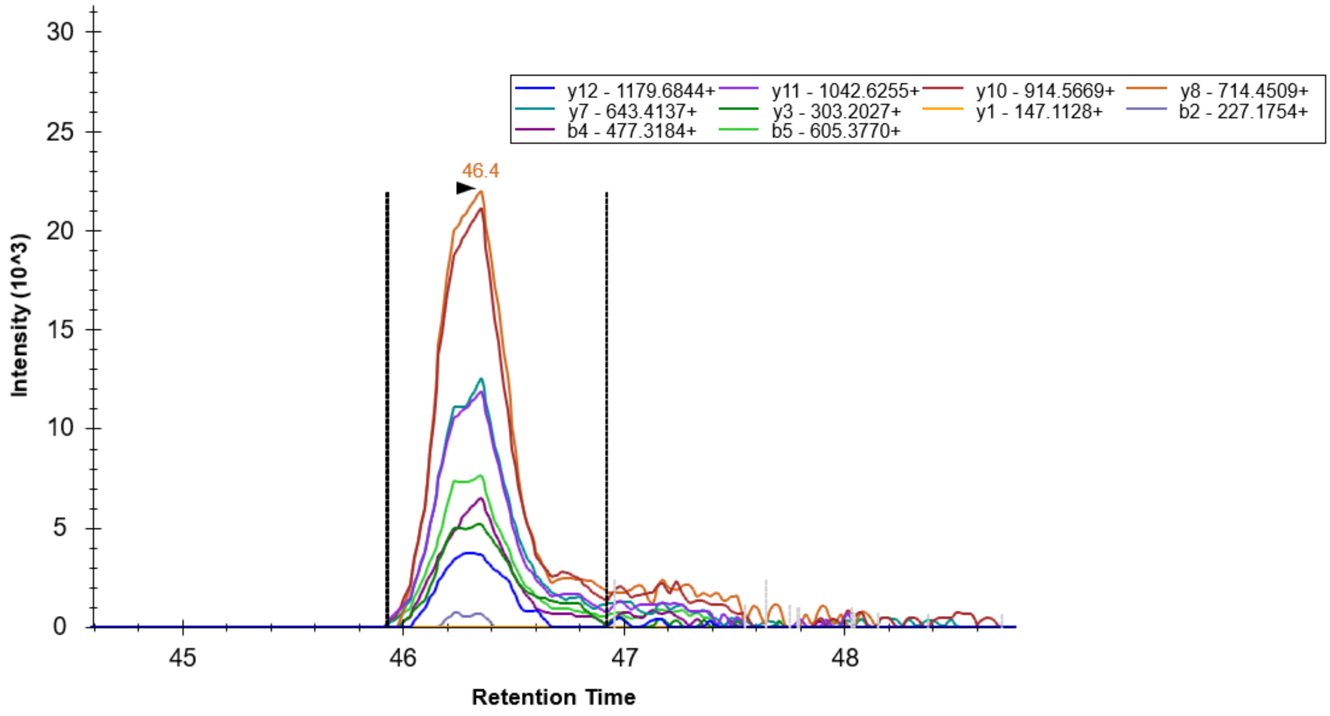


HNRNPK, TDYNASVSVPDSSGPER, 890.90284, 2+:

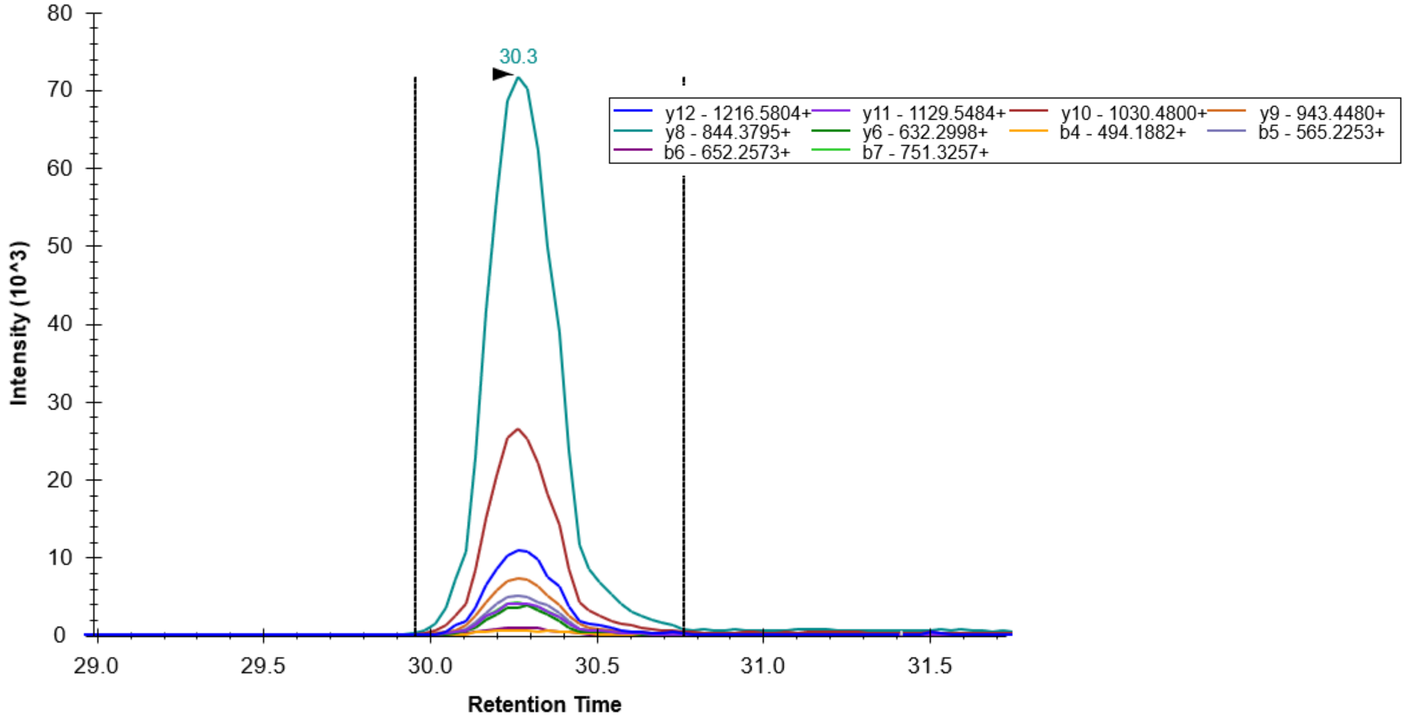

RPS16, LLEPVLLLGK, 547.86282, 2+

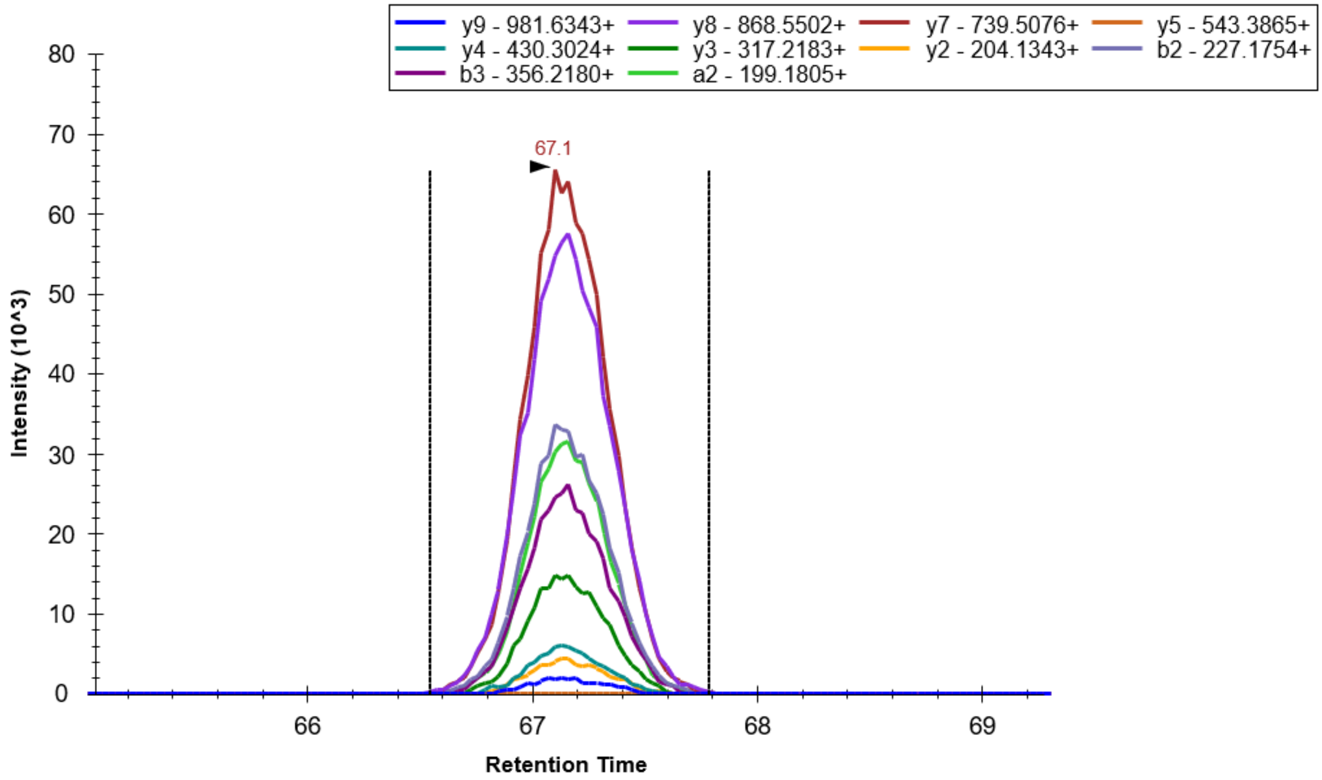


RPS16, GPLQSVQVFGR, 594.3302, 2+:

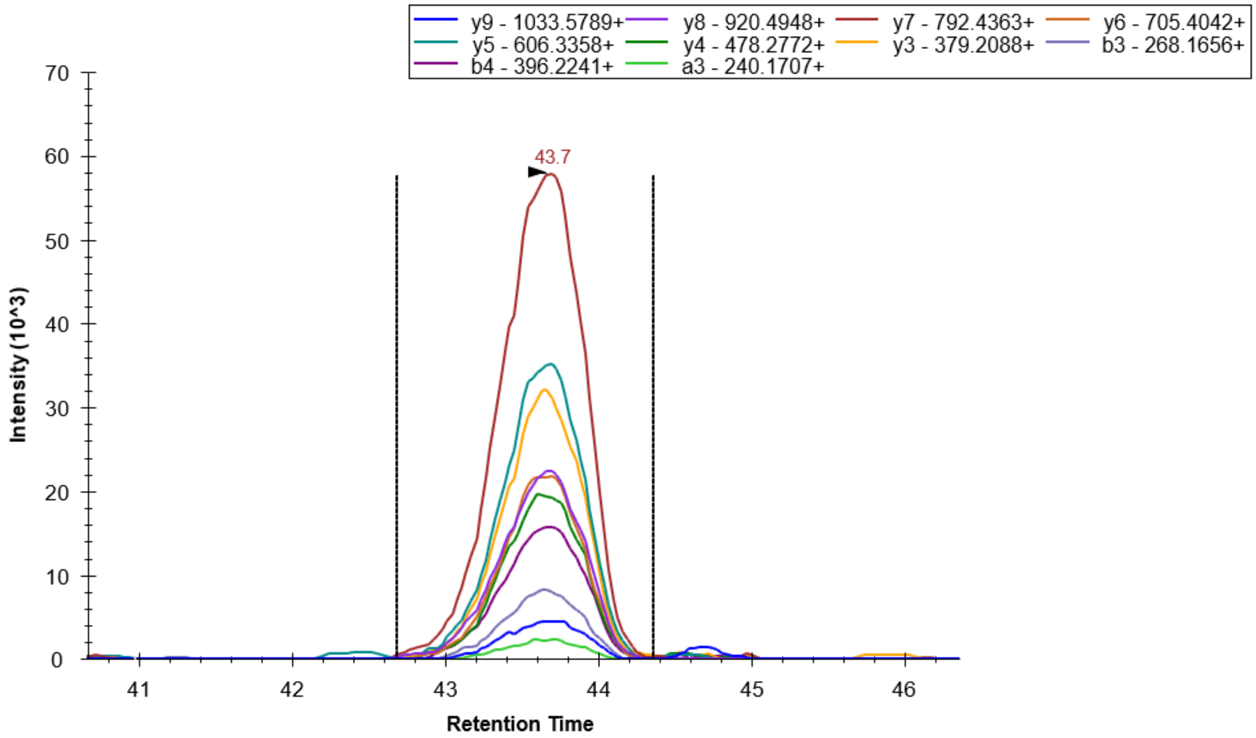

HNRNPA0, LFIGGLNVQTSESGLR, 845.95977, 2+:

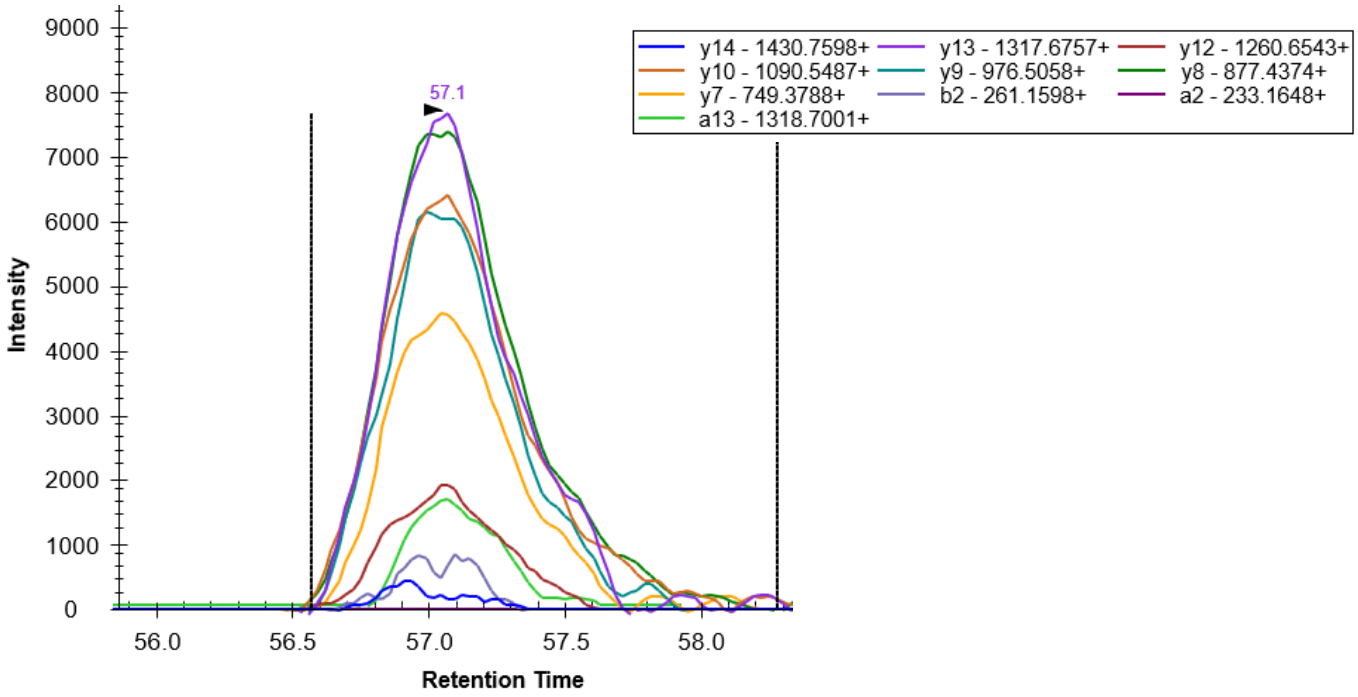


RACK1, VWQVTIGTR, 530.30092, 2+:

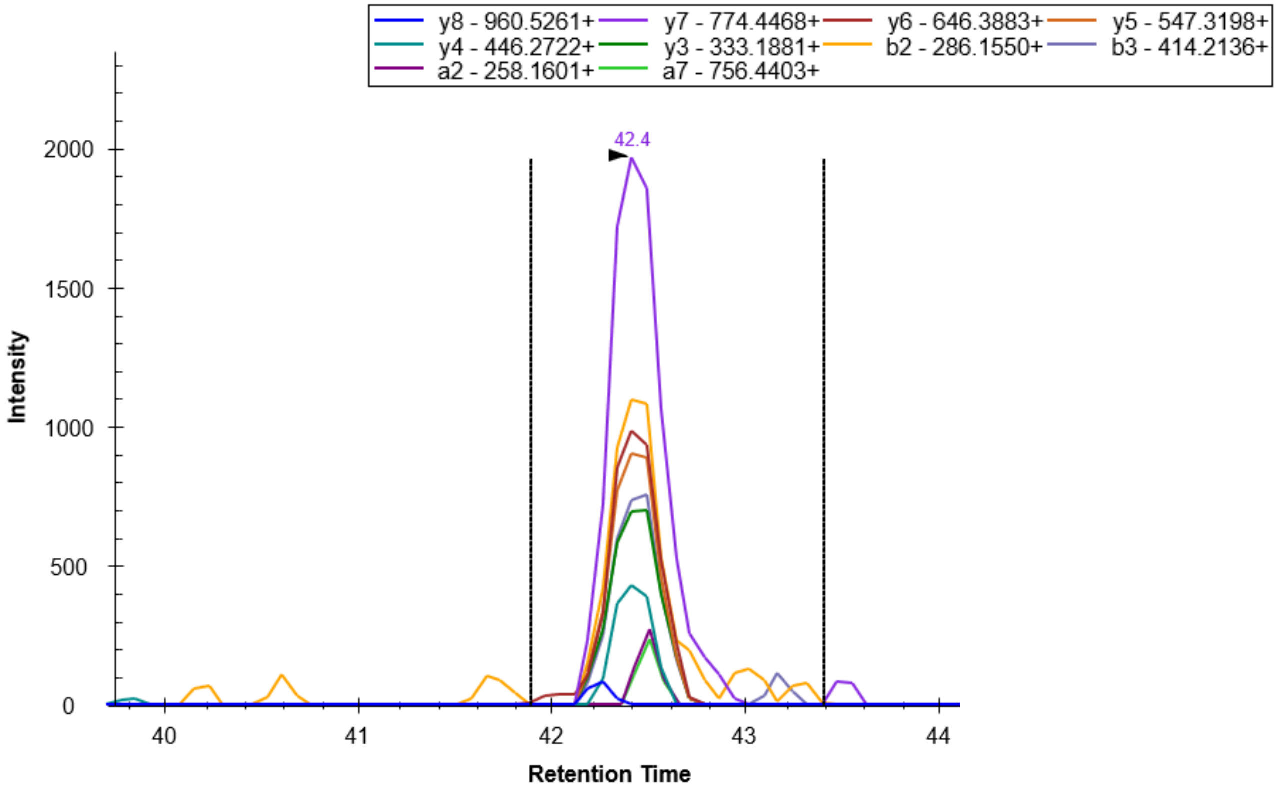

RACK1, DETNYGIPQR, 596.78328, 2+:

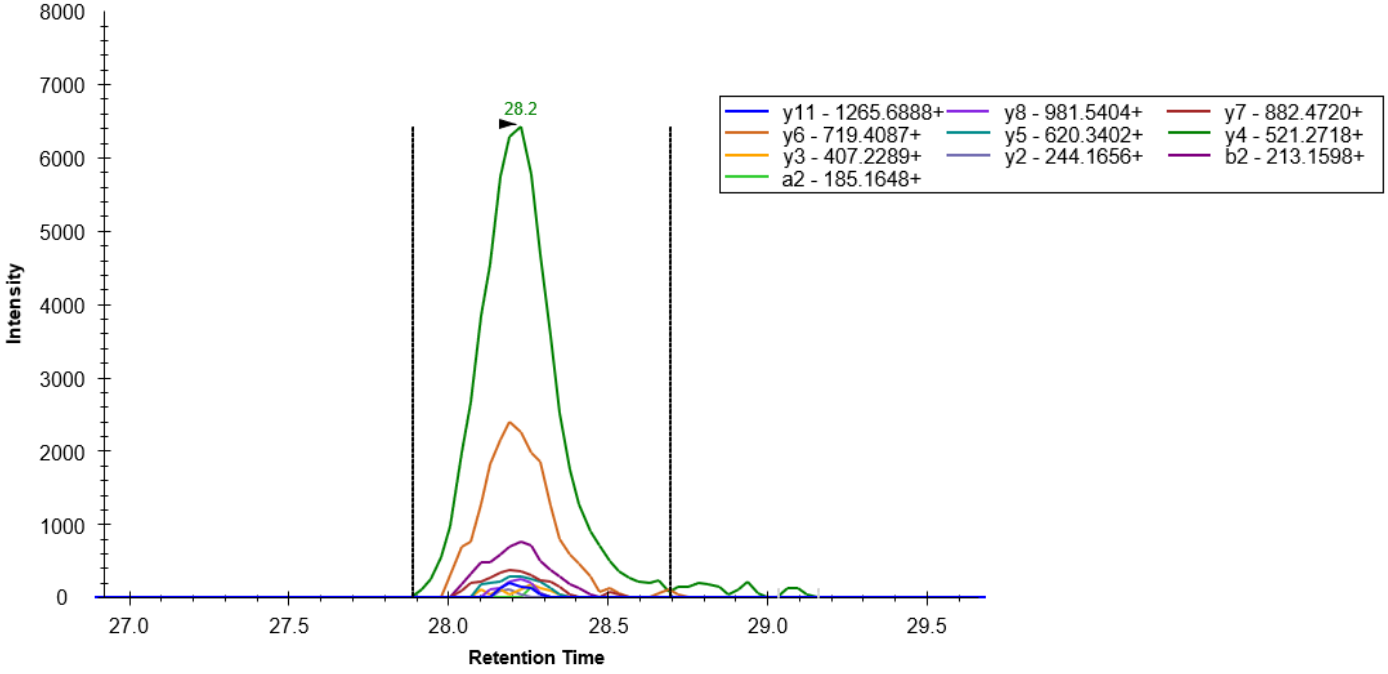


TARDBP, TSDLIVLGLPWK, 671.39247, 2+:

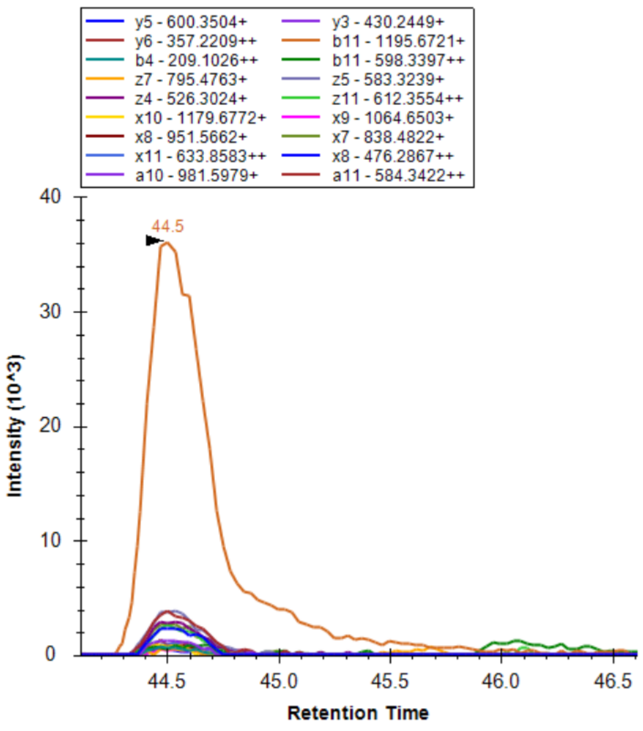

TARDBP, FGGNPGGFGNQGGFGNSR, 863.88767, 2+:

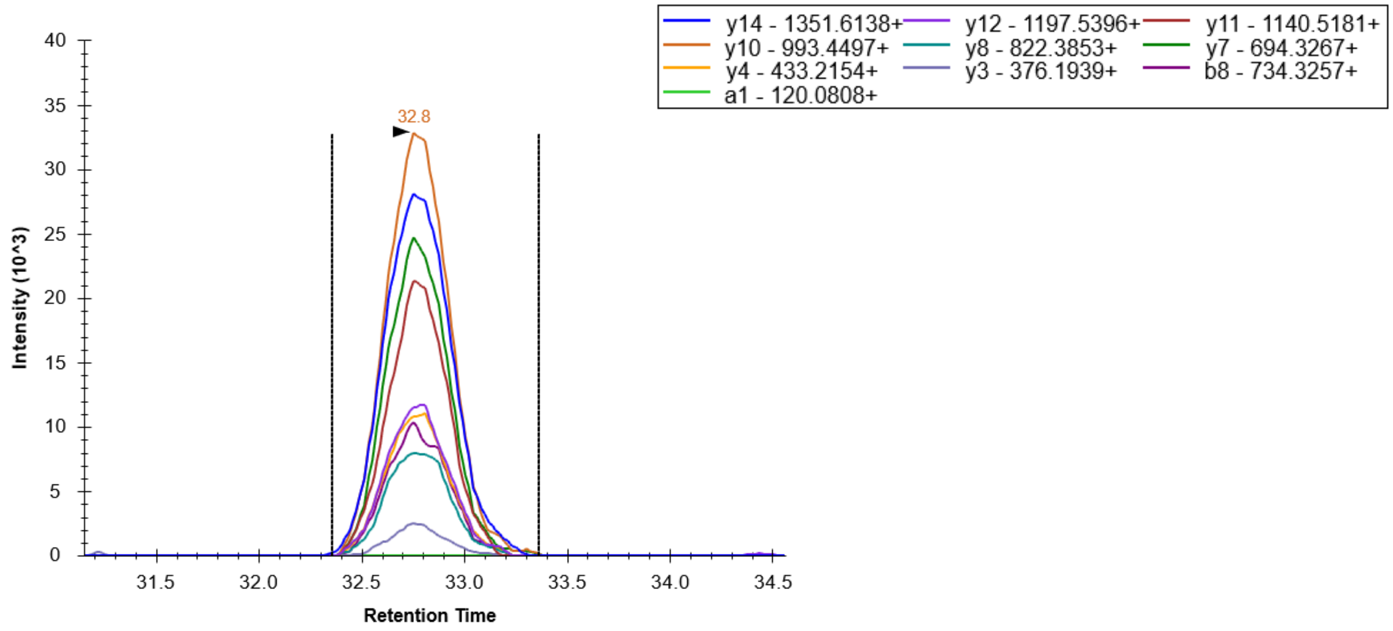


TARDBP, GISVHISNAEPK, 626.3382, 2+:

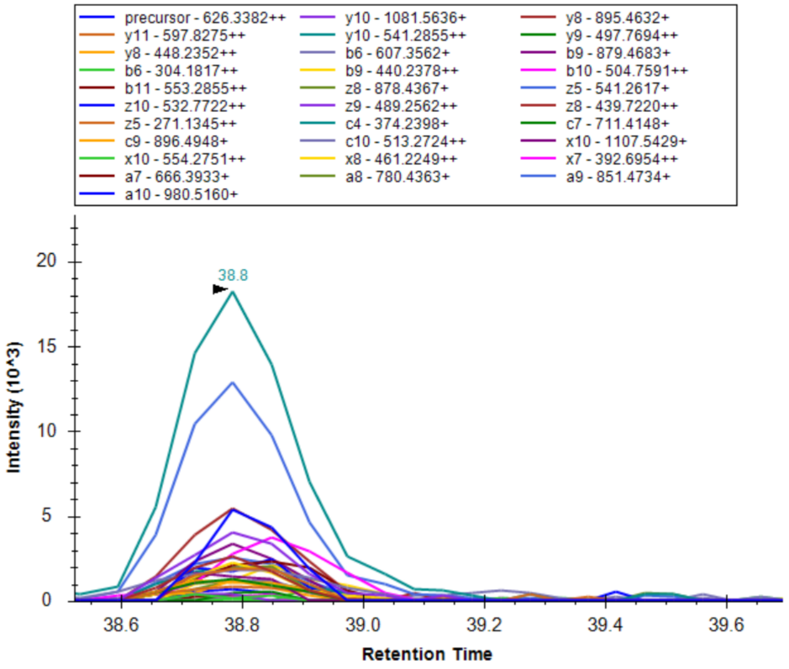

TARDBP, FTEYETQVK, 572.7797, 2+:

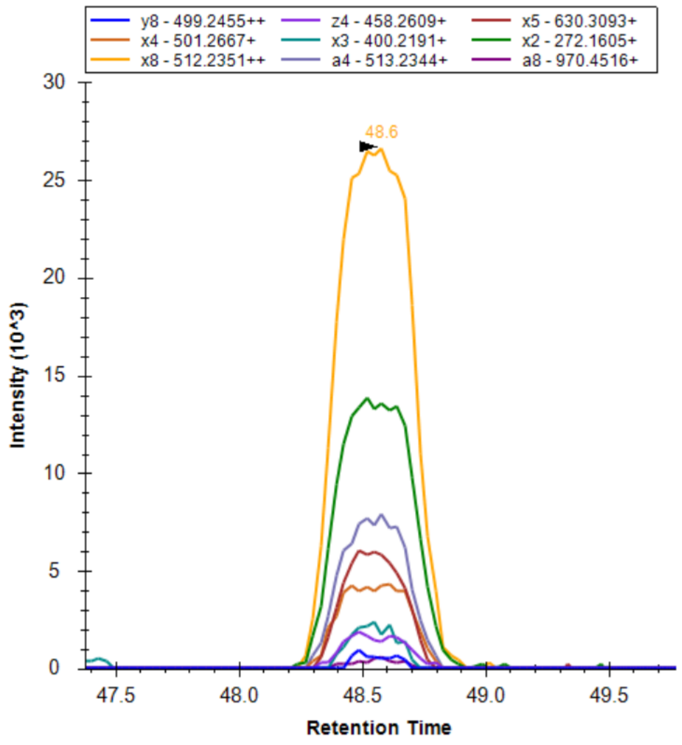


TARDBP, QSQDEPLR, 486.7409, 2+:

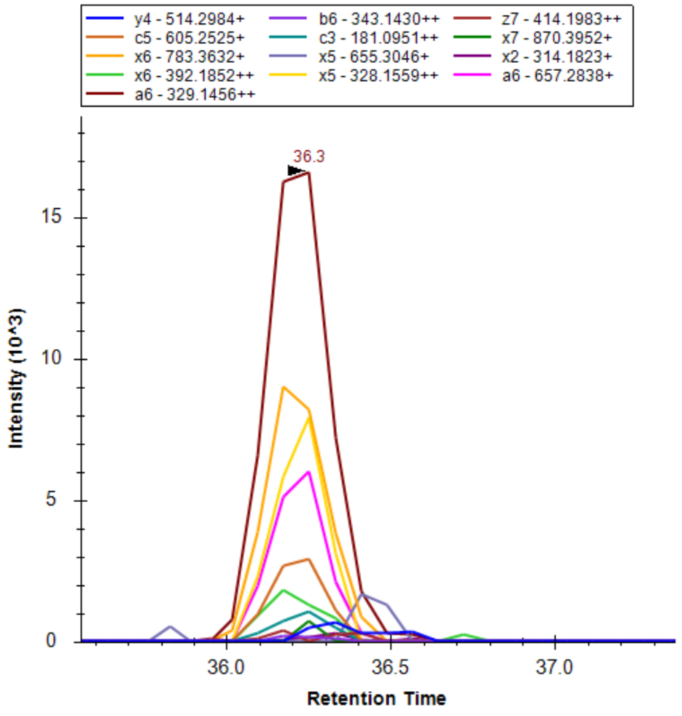

PDHA1, LEEGPPVTTVLTR, 706.3932, 2+:

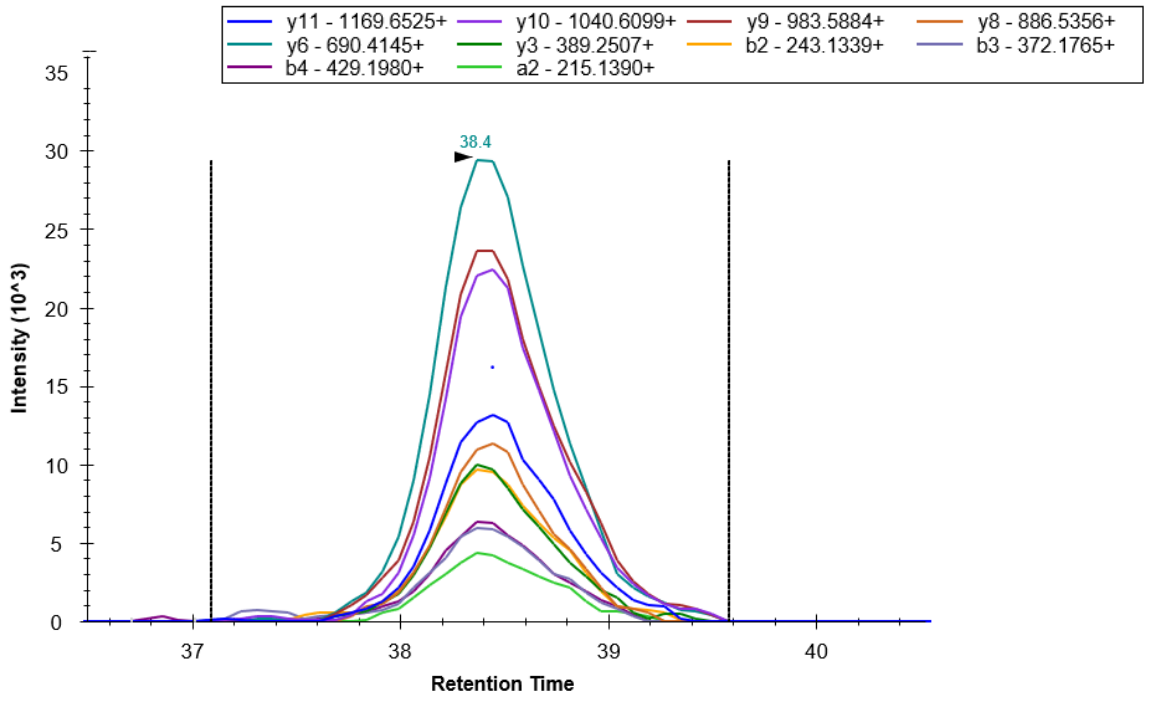


PDHA1, MVNSNLASVEELK, 717.3688, 2+:

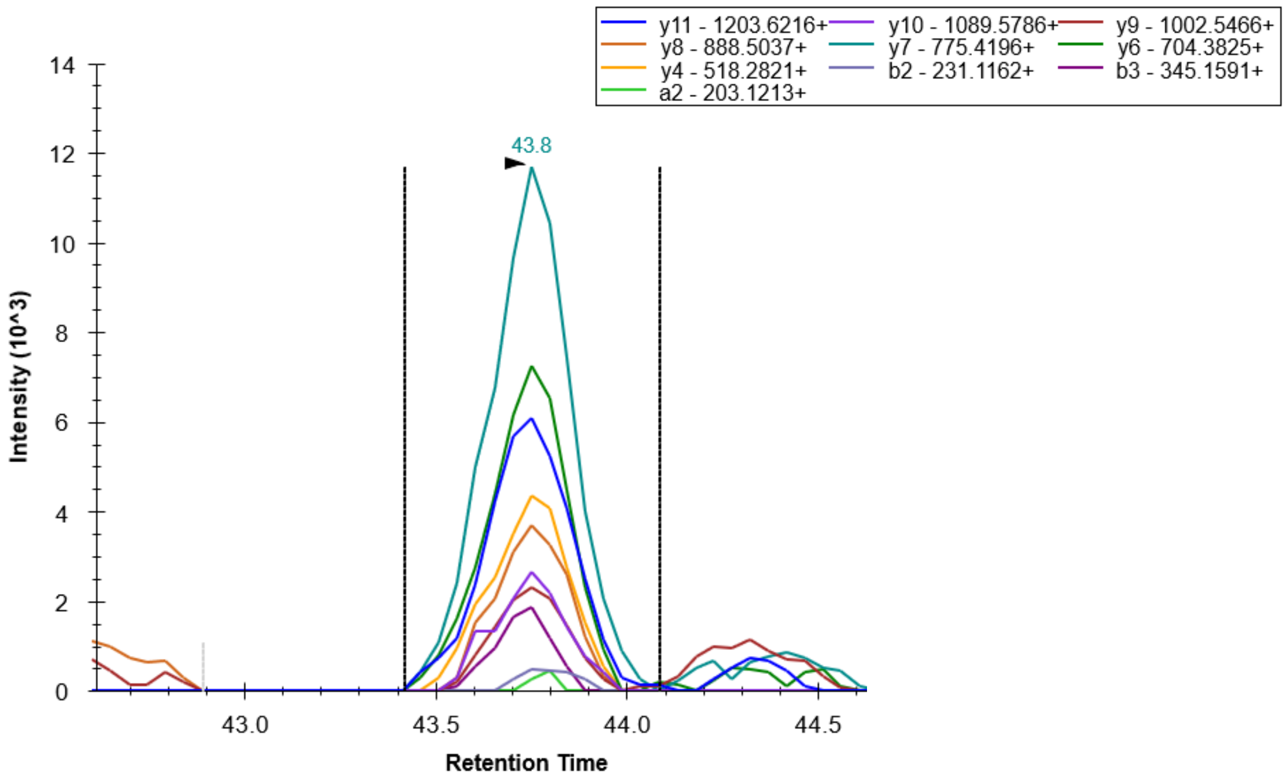

PDHA1, EILAELTGR, 501.2849, 2+:

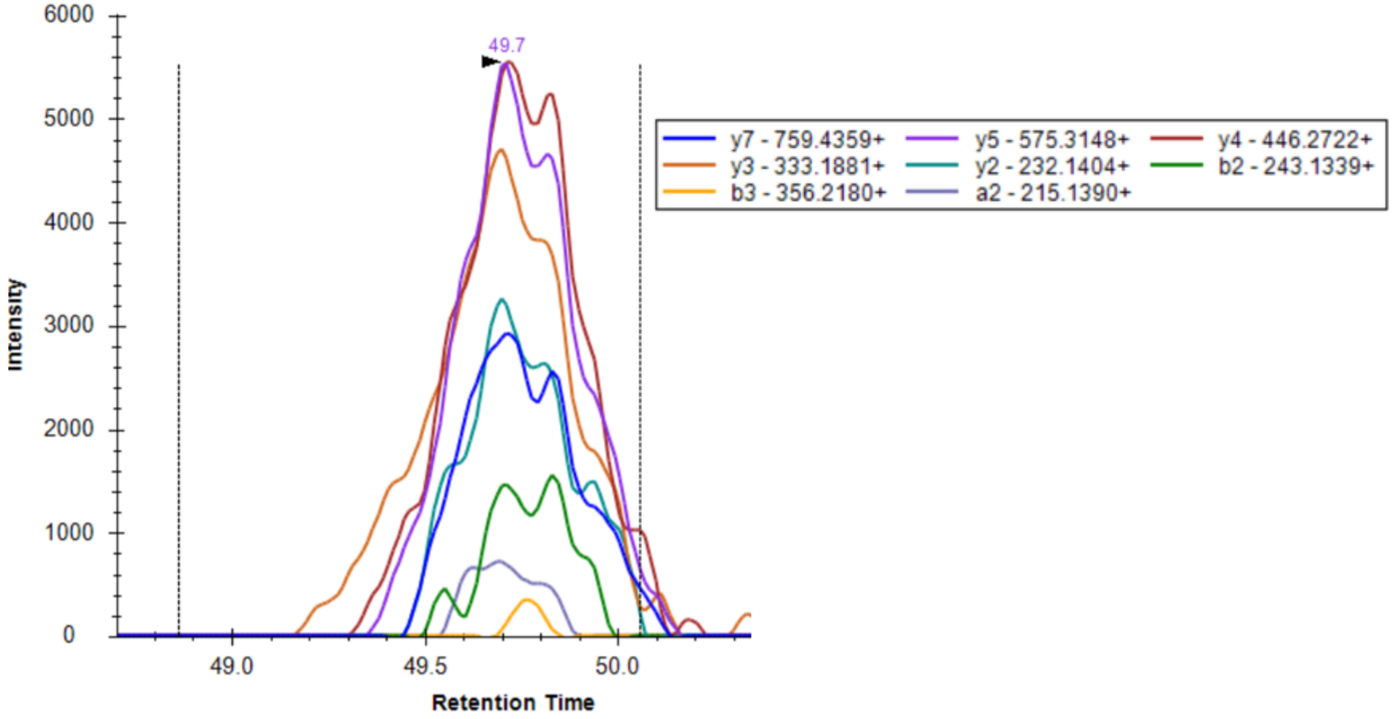


PDHA1, GPILMELQTYR, 660.8526, 2+:

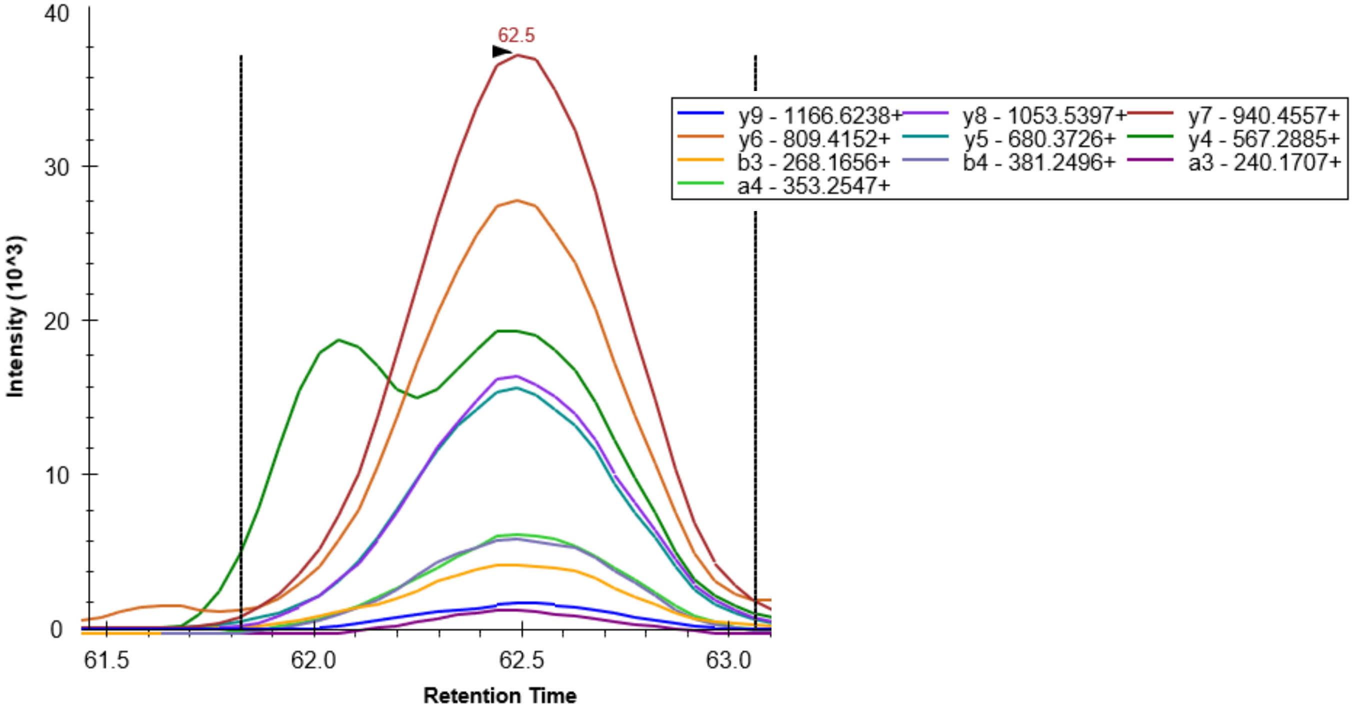

PDHB, TIRPMDMETIEASVMK, 926.4543, 2+:

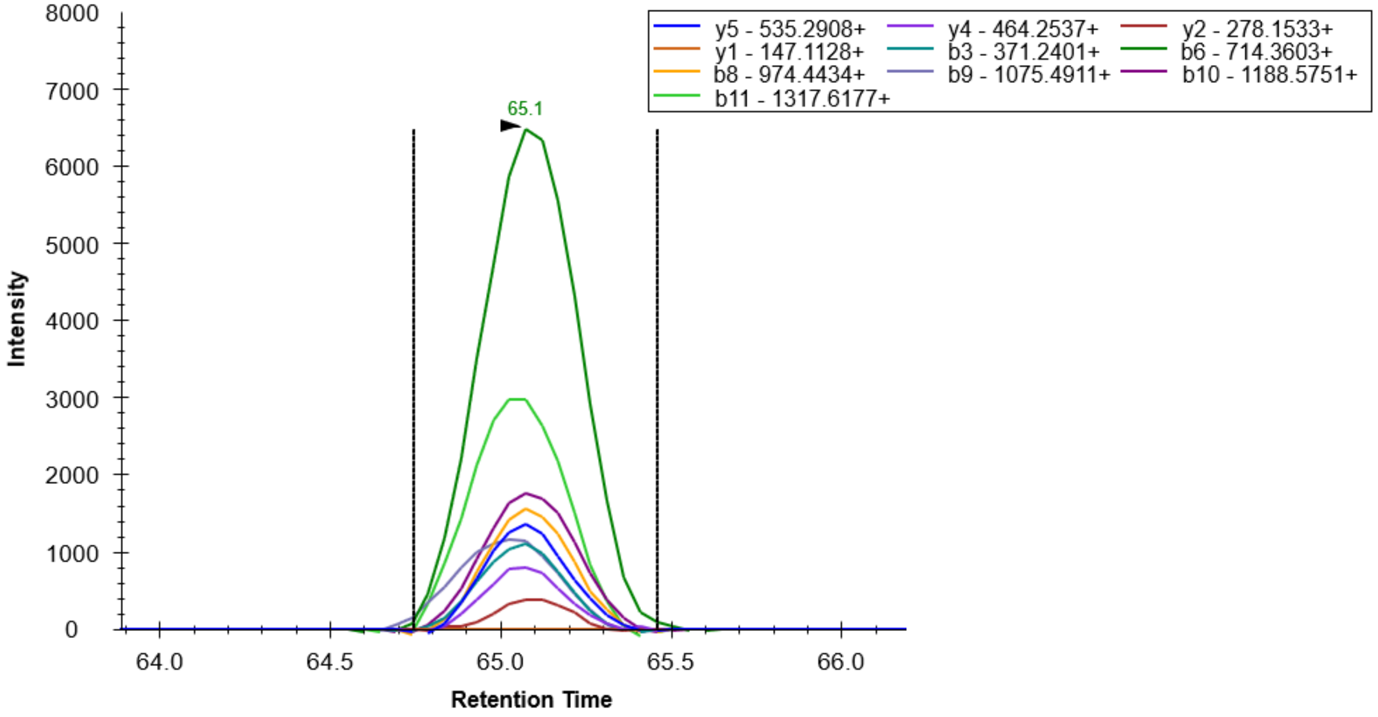


PDHB, VFLLGEEVAQYDGAYK,901.454, 2+

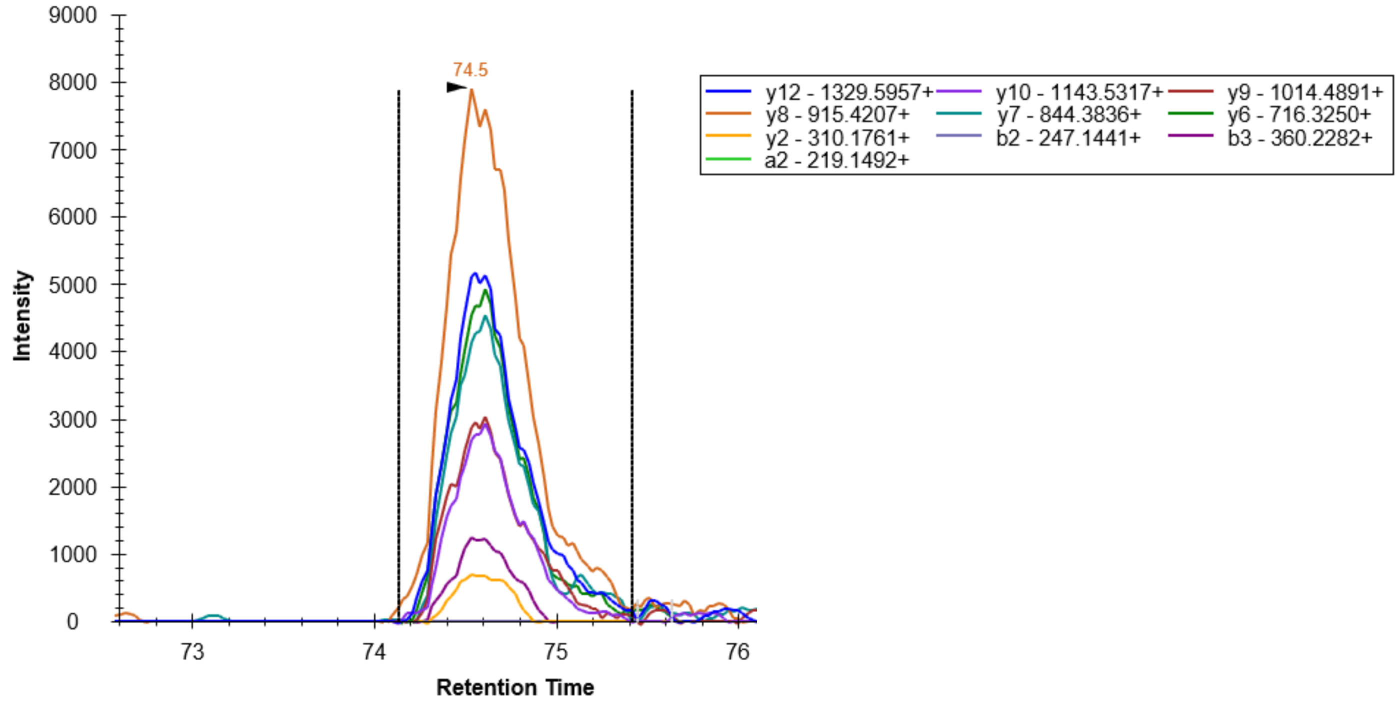

PDHB, IMEGPAFNFLDAPAVR, 874.4454, 2+:

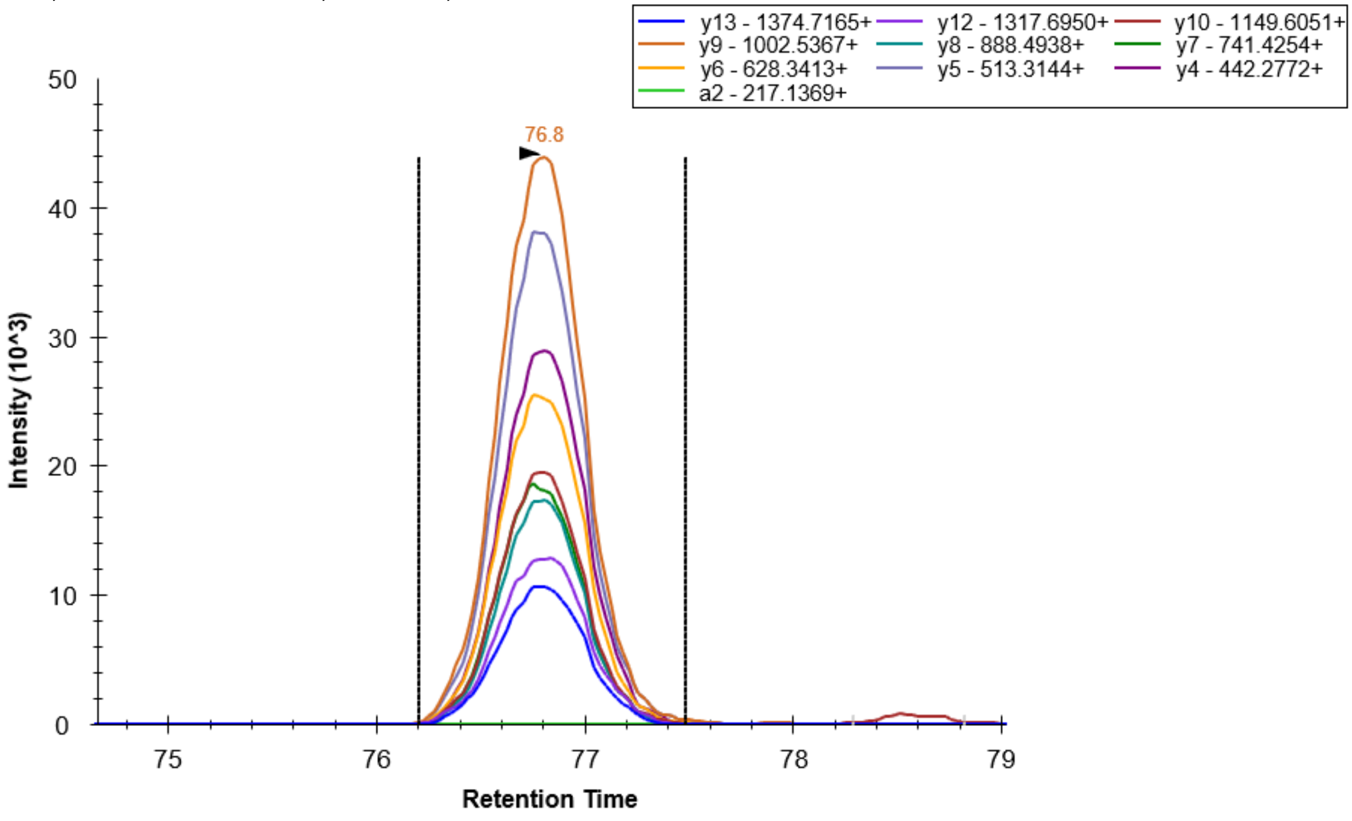


PDHB, DAINQGMDEELER, 760.3383, 2+:

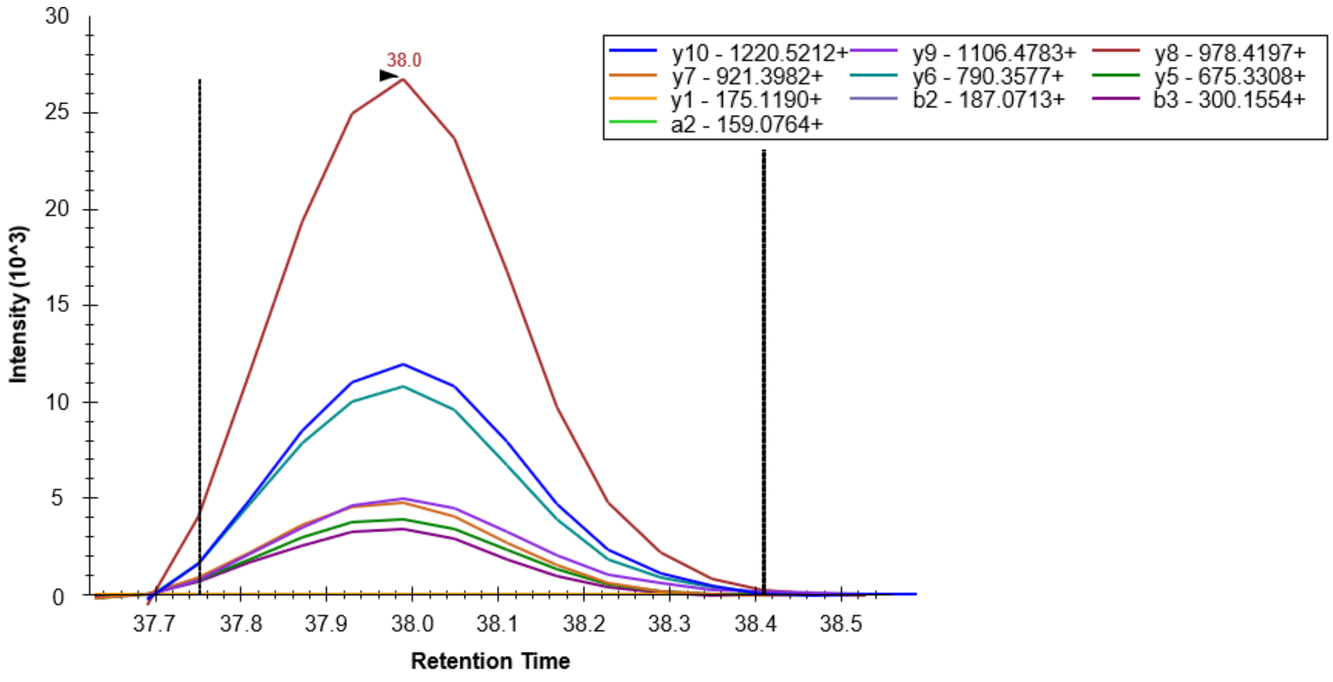

PDHB, ILEDNSIPQVK, 628.348, 2+:

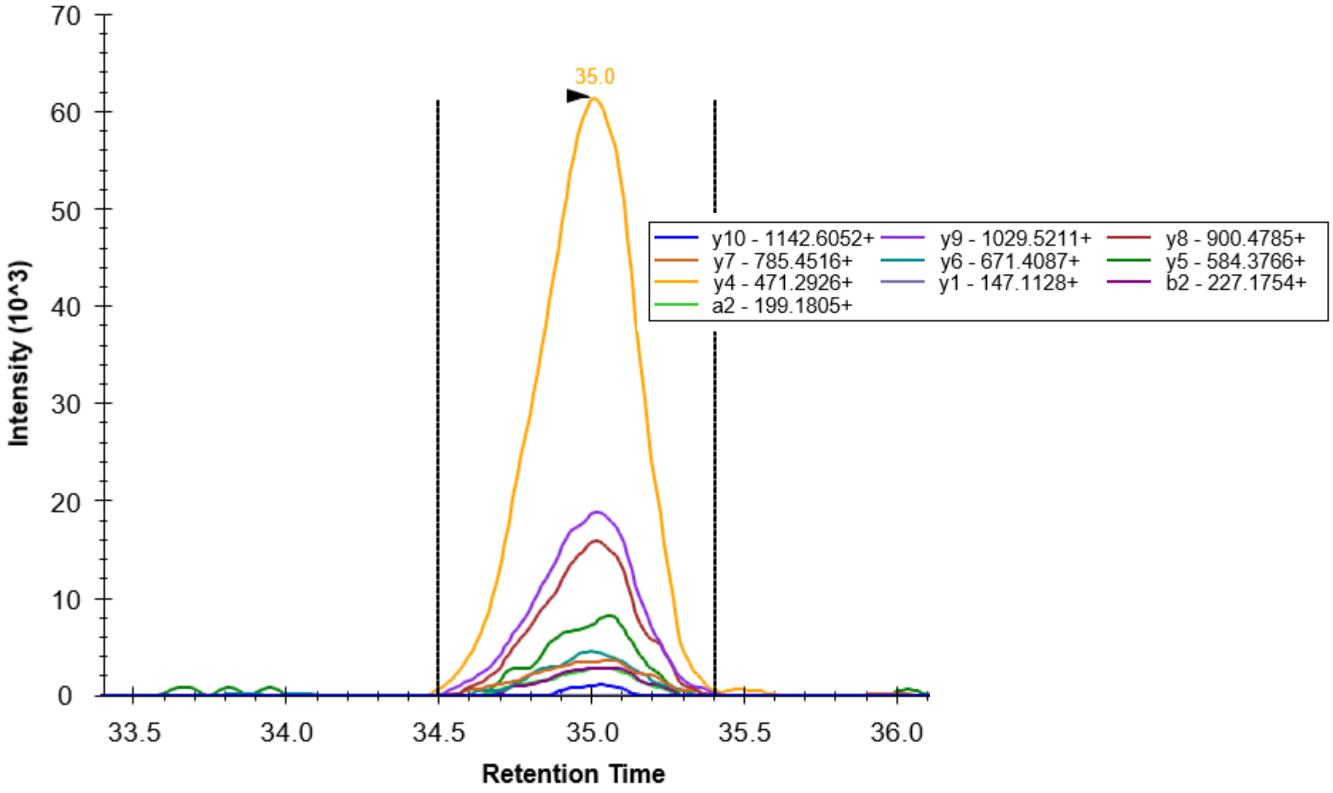


PDHB, DFLIPIGK, 451.7709, 2+:

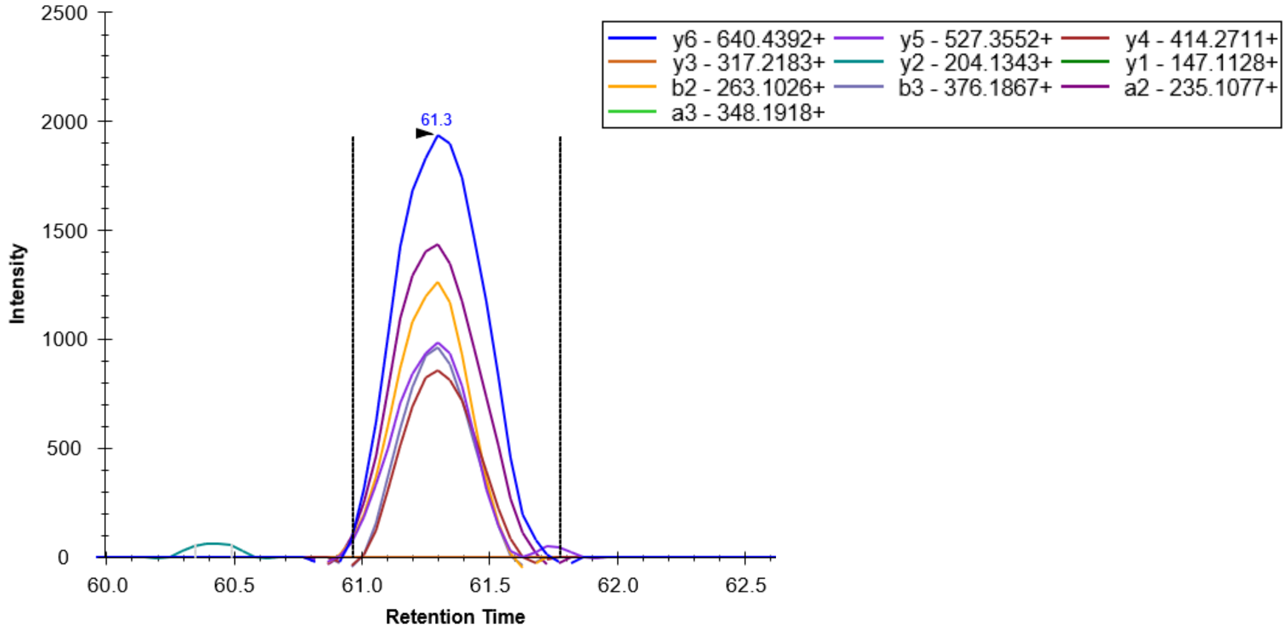

HNRNPK, 517.2223, 3+:

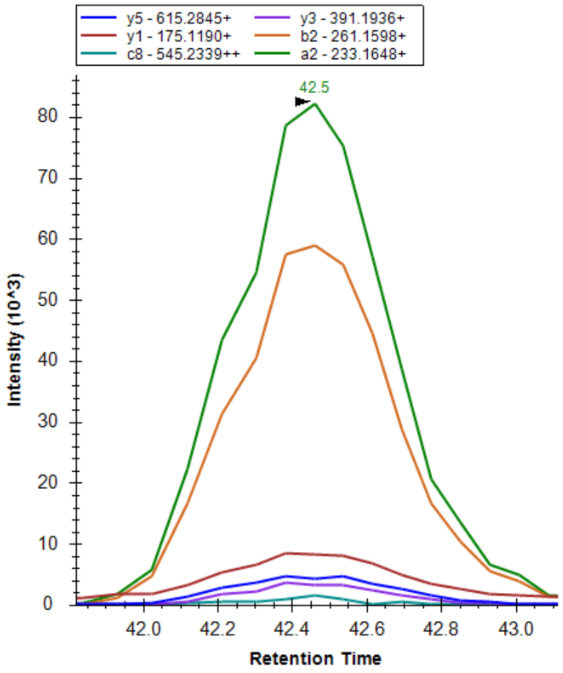


Internal Standard, VFFAEDVGSNK, 606.7984, 2+:

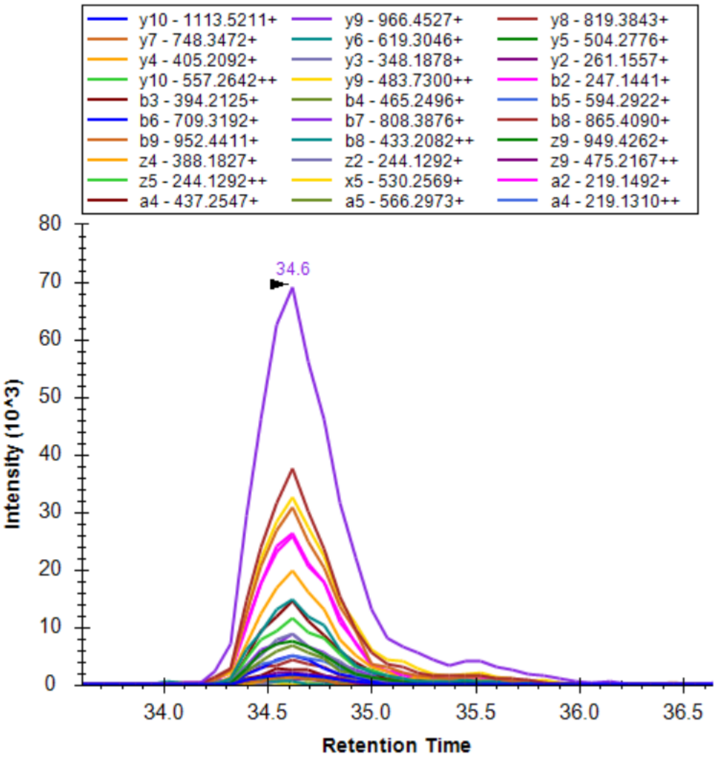
